# Practical quantification of immunohistochemistry antigen concentrations and reaction-diffusion parameters

**DOI:** 10.64898/2026.04.16.719078

**Authors:** Woody Perng, Berenice Mbiribindi, Benjamin Thomas Andrews, Xiangdan Wang, Debra Dunlap, Jeff Eastham, Hai Ngu, Andrei Chernyshev, Darya Orlova, Franklin V. Peale

**Affiliations:** Research Pathology, Genentech, Inc., South San Francisco, CA 94080, USA; Research Oncology, Genentech, Inc., South San Francisco, CA 94080, USA; BioAnalytical Sciences, Genentech, Inc., South San Francisco, CA 94080, USA; Voevodsky Institute of Chemical Kinetics and Combustion, Russian Academy of Sciences, Novosibirsk, Novosibirsk, 630090, Russia; Cancer Immunotherapy Discovery, Genentech, Inc., South San Francisco, CA 94080, USA

**Keywords:** Immunohistochemistry, reaction-diffusion, effective diffusion coefficient

## Abstract

The immunohistochemistry (IHC) methods widely used in diagnostic medicine and biomedical research are kinetically complex reaction-diffusion processes that, ideally, produce stain intensities correlated with the local antigen concentration. Yet after 75 years of use, practical theoretical tools to rigorously plan and interpret IHC experiments are still lacking. Because modeling the reactions requires time-consuming computer simulation, impractical for regular use, most protocols are optimized empirically, without detailed knowledge of the reaction rates and antigen-antibody equilibria. The resulting stain intensities can be calibrated against standards with known antigen abundance, but they are typically not interpretable in terms of chemical antigen concentrations. To address these limitations, we developed a fast interpolation method to model reaction-diffusion behavior, and experimental methods to characterize IHC kinetic parameters in formalin-fixed paraffin-embedded (FFPE) samples. Used together, these allow experimental measurement of both the chemical concentration of antigen in the sample and the reaction-diffusion parameters consistent with the assay results. Results show 1) direct immunofluorescent detection has low nanomolar sensitivity with >1000-fold dynamic range, and 2) antibody diffusion rates in FFPE samples can be >1000-fold slower than in aqueous solutions, producing diffusion-limited conditions in which the IHC reaction time course may depend on the sample antigen concentration. Awareness of these details is necessary to avoid potential underestimation of both the absolute and relative antigen concentrations in different samples that may occur if staining is stopped before reaching equilibrium. Software tools are provided to allow users to rapidly model IHC reaction time courses and to fit experimental time course data with candidate reaction parameters. The principles described here apply equally to other tissue-based “spatial omics” analyses and should be considered when designing and interpreting experiments requiring any macromolecule to diffuse into and react in a tissue section.

**SIGNIFICANCE:** The theoretical and experimental framework described here advances IHC staining from a qualitative or semi-quantitative method towards a more rigorously quantitative assay. The practical ability to predict IHC reaction kinetics and fit reaction parameters to experimental data has the potential to advance IHC applications in diagnostic medicine and biomedical research in three ways: 1) interpretation of experimental and diagnostic samples stained under different conditions can be more objective, facilitating comparison of results from different protocols and different laboratories; 2) IHC staining can be interpreted as molar chemical antigen-antibody concentrations calculated from the reaction parameters measured in the studied sample; 3) the correlation between antigen concentration and biological behavior can be examined more reliably. Practical software tools are provided.

## INTRODUCTION

Immunohistochemistry (IHC), immunofluorescence and related tissue-based assays are biochemical reaction-diffusion processes that create spatially precise signals reflecting the relative abundance of the target antigen. Typical samples for staining on a glass slide are 4-10 µm-thick sections cut from formalin-fixed, paraffin-embedded (FFPE) or frozen specimens, but thicker samples may be stained and imaged with “deep IHC” methods (1,2). While IHC was initially used as a qualitative molecule-specific “special stain’’ to detect diagnostically informative antigens, it is increasingly evaluated more quantitatively to inform patient prognosis and therapeutic decision-making (3,4). However, a practicable theoretical framework to guide IHC experimental planning and interpretation is lacking, making the quantitative evaluation of results challenging. Our goal in this work was to address this need.

The endpoint of an IHC assay is most often a binary presence/ absence or semiquantitative assessment of antigen. However, as is widely appreciated, quantification of IHC results is complicated by the large number of assay variables including sample fixation, antigen retrieval, antibody binding conditions, and chromogenic or fluorescent detection methods. There are no universal standard assay conditions: representative “ready-to-use” antibody on-slide concentrations vary more than 150-fold (Table S1); antibody affinities measured *in vitro* vary more than 1000-fold (5) and primary antibody incubation times can range from minutes to days (6,7). The limit of detection and dynamic range, which should correspond to biologically meaningful antigen levels, may vary between empirically optimized assays. Assay sensitivity can be adjusted over a wide range; similar-appearing strongly positive chromogenic signals may correspond to antigen copies per cell that vary more than 5,000-fold, from 2*10^6^ [HER2 in SK-BR-3 cells (8)] to 10^10^ [insulin in pancreatic islet β-cells (9)].

The practical consequences of IHC assay variability are clinically relevant: comparison of results from three FDA-approved PD-L1 assays is complicated by differences in their sensitivity and dynamic range (10–13), and similar clone-dependent variations in antigen detection have been documented for HER2, EGFR, ER and ALK1 (14–17). More recently, antibody-drug conjugates with activity in “HER2-low” patients have made necessary the reproducible evaluation of HER2 even at the threshold of detectability (18). Unfortunately, the diversity of possible assay conditions means that without a detailed understanding of IHC reaction kinetics, stain intensity cannot be interpreted to represent either an absolute antigen concentration or a known fraction of the total detectable antigen in the tissue sample.

A 4 µm-thick tissue section is ∼300 times thicker than the 12-15 nm diameter of an IgG molecule, requiring the antibody to diffuse into the sample to reach all the available antigen. To accurately measure and model IHC reaction behavior, the diffusion rate of antibody through the section must be quantified and compared with the antigen-antibody reaction rate. While it is widely recognized that diffusion can be rate-limiting in thick samples (1), it is unclear whether this is true in the thinner samples typical of diagnostic IHC practice. Despite early evidence that diffusion limitations are relevant even in cell monolayers (19), attention to this issue has been inconsistent (20–23). When antibody diffusion rates have been modeled in IHC, the values are often taken from experiments in living tissue or aqueous buffers rather than being measured in the sample analyzed (22,24). The potential errors are large because the effective diffusion coefficient of IgG in buffer is ∼4.0-4.4E-7 cm^2^ sec^-1^ (25,26), while the reported rates in unfixed living tissue are from 3 to >50 times slower (26–29), and diffusion rates for IgGs in fixed tissues have not been reported.

This work addresses three challenges to resolving these issues. First, accurate planning and interpretation of IHC experiments requires knowledge of the target antigen concentration ([Ag]), the antibody effective diffusion coefficient (D_eff_), and the forward and reverse reaction rate constants (k_for_ and k_rev_), all measured in the conditions used to stain the experimental sample. Second, practical analysis tools are needed to predict IHC reaction behavior and to fit candidate reaction parameters to experimental data. Third, while the progress of solution-phase Ag-Ab reactions can be described algebraically, the fact that IHC assays are hybrid reaction-diffusion processes complicates mathematical analysis. As is standard in the theory of nonlinear partial differential equations (30), there exists no general method to derive closed-form analytical solutions for non-integrable systems such as the general reaction-diffusion equation. Instead, computer simulation is required for each set of conditions. But as has been noted (31), even if the necessary computational resources and expertise are available, it is impractical to systematically search through a multi-parameter space to find conditions that meet the goals of the intended use, or fit experimental data.

A framework to solve these problems exists in established chemical engineering and surface plasmon resonance (SPR) theories that quantify the relative time scales for the separate reaction and convection/diffusion processes. In chemical engineering, the efficiency of porous metal catalyst particles can be characterized by a unitless “effectiveness factor”, *eta* (η; 0 < *eta* ≤ 1) which, under hypothetical quasi-steady-state conditions, defines the average catalytic activity of the entire particle volume relative to the activity at the external surface (32). Similarly, in SPR theory, a “thickness factor” mathematically identical to *eta* quantifies diffusion-limited reaction behavior in the ligand-containing hydrogel layer on an SPR chip surface (33). Another unitless parameter, a form of Damköhler number (Da), quantifies the reaction rate on the SPR chip surface relative to the rate of reactant delivery by convection and diffusion (31). However, important differences between SPR and IHC experimental geometries and procedures require that different analytical approaches be used. We show below that three parameters, *eta*, Da, and the ratio of antibody concentration to the antigen-antibody dissociation constant ([Ab]/K_D_), usefully inform IHC reaction analysis.

The imaging and analytical methods described here allow objective quantification of the molar concentration of antigen and estimation of the Ag-Ab reaction kinetic parameters in fixed samples. A software tool allowing rapid prediction of IHC kinetics is used to objectively define candidate reaction parameters consistent with experimental data. Notably, the ability to measure tissue antigens in absolute molar concentrations rather than as relative intensities should make it possible to more accurately compare results from IHC assays done under different conditions, in different labs, and to more accurately correlate antigen expression with biological behavior. More broadly, the general concepts presented here are relevant to multiplexed spatial “omics” analyses, expansion microscopy, immunotherapy for solid tumors and other situations where diffusion-limited binding reactions occur.

## MATERIAL AND METHODS

### Analytical solution for diffusion in a plane sheet

As a reference against which to compare diffusion rates modeled in software, the rate of diffusion from a source of constant concentration into a finite volume lacking specific and non-specific binding sites was calculated according to Eq. 2.54 from (34):

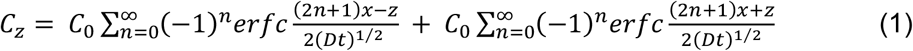

(Mathematical abbreviations are summarized in Table 1.)

**TABLE 1.** Mathematical abbreviations.

| Abbreviation | Meaning | Units |
| --- | --- | --- |
| [Ab] | Antibody concentration | M |
| [Ag] | Antigen concentration | M |
| [Ag]<sub>o</sub> | Original antigen concentration | M |
| [Ag]<sub>t</sub> | Antigen concentration at time *t* | M |
| [AgAb] | Concentration of AgAb | M |
| [AgAb]<sub>equil</sub> | Equilibrium AgAb concentration | M |
| C<sub>z</sub> | Concentration at point z | M |
| C<sub>o</sub> | Original concentration | M |
| C*r<sub>f</sub>* | Crank constant at *f* | none |
| D | Diffusion coefficient | μm<sup>2</sup>s<sup>-1</sup> |
| D<sub>eff</sub> | Effective diffusion coefficient | μm<sup>2</sup>s<sup>-1</sup> |
| *erfc* | Complementary error function | none |
| *f* | Fractional reaction completion, *f* | none |
| K<sub>D</sub> | Dissociation constant | M |
| k<sub>eff</sub> | Effective reaction rate constant | s<sup>-1</sup> |
| k<sub>for</sub> | Forward reaction rate constant | M<sup>-1</sup>s<sup>-1</sup> |
| k<sub>rev</sub> | Reverse reaction rate constant | s<sup>-1</sup> |
| *t* | Time | s |
| τ<sub>D</sub> | Characteristic diffusion time | s |
| τ<sub>Rx</sub> | Characteristic reaction time | s |
| x | Sample thickness | μm |
| z | Distance from glass slide surface | μm |

The equation was solved for values of *z / x* = 0 to 1 in 1000 steps, summed over n = 0 to 10, and the integrated area under the resulting curve was estimated by trapezoidal approximation.

### Analytical description of solution-phase AgAb reactions

For a solution-phase Ag-Ab binding reaction, assuming a constant concentration of bivalent Ab (*i*.*e*., Ab having 2 antigen-binding Fab domains), the equilibrium AgAb concentration relative to the original antigen concentration is:

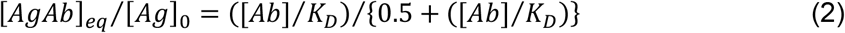

Under the same conditions, fractional progress towards reaction equilibrium, *f*, at any given time *t* (in seconds) is

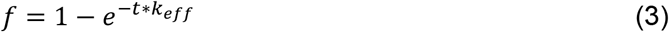

where

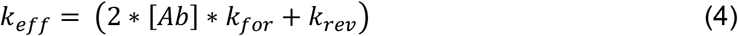

and the time, *t*, required to reach a given fractional reaction completion is

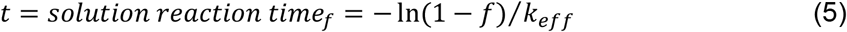

### Characteristic times for diffusion and AgAb binding reaction

The characteristic diffusion time (s), *τ*_*D*_, and reaction time (s), *τ*_*Rx*_, are

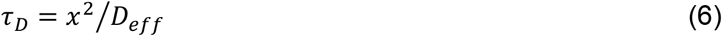

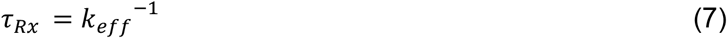

Restating Eq. 5 in terms of *τ*_*Rx*_ gives:

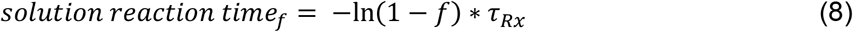

“Null diffusion time_*f*_” is the time needed for the average [Ab] in a volume unreactive with antibody to reach a given fraction of the equilibrium final value, *f* = [Ab]_t_ / [Ab]_0_:

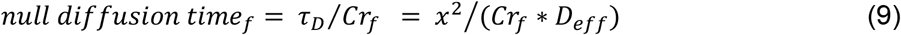

Representative values of *Cr*_*f*_, obtained from Eqs. 1 and 9, are given in Table S2.

To relate the solution-phase reaction times with the corresponding IHC reaction times, define *Iota*_*f*_ = solution-phase reaction time_*f*_ / IHC reaction time_*f*_

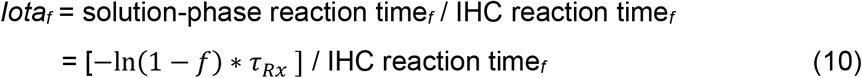

### Kappa relates τ_D_ **and** τ_Rx_

To facilitate modeling and data analysis for reactions with different values of *τ*_*Rx*_ and *τ*_*D*_, we defined *kappa*_*f*_, a form of Damköhler number, as the null diffusion time relative to the solution reaction time when each is evaluated at a given fractional progress, *f*, towards reaction equilibrium:

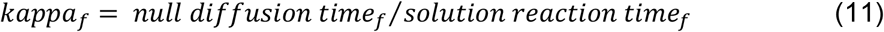

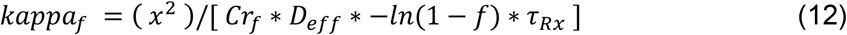

### *Eta* quantifies relative tissue [Ab] early in the IHC reaction

Thiele’s definition of the “effectiveness factor”, *eta*, for a first-order irreversible chemical reaction occurring with no change in volume in a plate-shaped catalyst particle (32) was adapted for IHC reactions in sections of uniform thickness as:

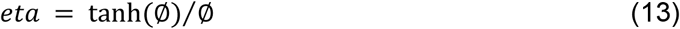

where:

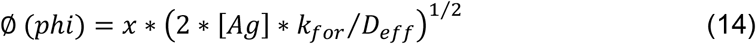

*Eta* (0 < *eta* ≤ 1) quantifies the average [Ab_free_] in the sample volume relative to the solution-phase [Ab] at a quasi-steady-state condition early in the IHC reaction when the reverse reaction is negligible and free antigen has not yet been significantly converted to AgAb product. As the value of *eta* decreases, the initial average [Ab_free_] in the sample decreases, and the IHC reaction therefore proceeds more slowly than the corresponding solution-phase reaction. *Eta* < 0.95 when ∅ > 0.4.

### Virtual Cell and MATLAB reaction-diffusion models

IHC reaction time courses were simulated using Virtual Cell (VCell) 7.7.0 (35,36) and a custom program written in MATLAB [version 9.10.0 (R2021a) Update 3; The MathWorks Inc., Natick, Massachusetts]. VCell two-dimensional models typically had a 1 µm-deep layer of antibody solution at a fixed concentration overlying a 4 µm-thick layer of uniform [Ag]. Simulations used the fully-implicit finite volume, regular grid (variable time step) algorithm, with absolute and relative error tolerances of 1E-9 and 1E-7 respectively, 0.005-0.1 um grid dimensions in all axes, and 1E-4-1E-2 s time steps. “Disable Border Extrapolation” was not selected. Detailed models are available for download at GitHub (https://github.com/Genentech/IHC-reaction-diffusion).

The alternative MATLAB reaction-diffusion model is based on the concept that diffusion is a stochastic process in which a constant fraction of the mobile molecules in a unit volume move to the adjacent unit volumes in each unit of time, regardless of the local concentrations. The macroscopic result is the time-dependent equalization of concentrations, but the motion of individual molecules is independent of concentration gradients (34). Based on this principle, the model simulates diffusion algebraically by dividing the sample thickness into a selectable number of layers and moving, at each time step, a fixed fraction (10% by default) of the free antibody concentration in each tissue layer to each of the immediately adjacent tissue layers. The model time steps are calibrated to reproduce diffusion rates in the absence of Ag (Eq. 1). The length of a timestep (in seconds) is:

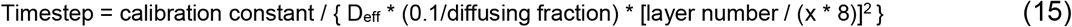

With default values for diffusion fraction (10%), D_eff_ (0.25 µm^2^s^-1^), tissue thickness (x = 4 µm), and layer number (8/µm of tissue thickness), the calibration constant is 1.55041E-3 and the corresponding time step is 3.876025E-4 seconds. The time step is recalculated by the software when D_eff_, diffusing fraction, layer number or tissue thickness are changed. Calculations (Eq. 1) show that a 10% diffusing fraction is an acceptable approximation. In one time step, <1% of this amount (*i*.*e*., < 0.1%) would move beyond the immediately adjacent unit volume in a real-world diffusion process. Choosing a smaller diffusing fraction (less than 10%) reduces this error at the cost of increasing calculation time.

The model calculates for each time step the forward and reverse Ag-Ab reactions using the local reactant concentrations in each layer, according to irreversible second-order forward and first-order reverse reaction kinetics. Simulations were run for fractional reaction completion *f* = 0-0.9 for 2294 parameter combinations systematically distributed from log_10_([Ab]/K_D_) = -2 to 5, log_10_(*kappa*_*f*_) = -3 to 2, and -log_10_(*eta*) = 0.02-1.6. Results are recorded at user-selected time points. Average sample reactant concentrations in the sample volume are estimated by trapezoidal interpolation.

### Reagents

Buffer components including bovine serum albumin (cat. # A5611-10G), ProClin-300; propyl gallate, Tween-20 were from MilliporeSigma, reagent grade unless noted otherwise. Other reagents and their sources were: anti-BCL2-AlexaFluor™-647 (cat. # NBP2-34513AF647) and mouse IgG isotype control clone 11711 AlexaFluor™-647 (cat. #: IC002R; Novus Biologicals, LLC); normal goat serum (cat. # 005-000-121; Jackson ImmunoResearch Laboratories); bovine serum albumin (BSA)-AlexaFluor™-647 (cat. # A34785; ThermoFisher). Flow cytometry reagents and sources were: Cytofix/Cytoperm (cat. #: 554714; BD Biosciences); Fixable viability dye eFluor 450 (cat. #: 65-0863-14; Invitrogen); Human TruStain FcX (cat. #: 422302; BioLegend).

### Sample preparation

Tissue microarrays: Synthetic antigen control samples containing BSA (12.5% final concentration) and 2.5E-8 M to 2.5E-4 M peptide encoding human BCL2 (UniProt P10415) a.a. 41-54 with N- and C-terminal extensions of Ac-YGSG and GSGC-amide, respectively, were prepared as described previously (37). Packed cell pellets were prepared from RAMOS (negative control; BCL2 nRPKM =0.15) and Granta-519 (positive control; BCL2 nRPKM =101) cultures, fixed in neutral-buffered formalin and embedded in paraffin using standard tissue processing methods. A tissue microarray was constructed using triplicate 1 mm diameter cores punched from donor blocks, arrayed in a regular 7 x 7 grid using a TMA Grand Master automated arrayer (3DHistech, Budapest, Hungary.)

Coverslip preparation: 22.5 mm square #1.5 thickness (0.17 mm) coverslips (cat. # 72204-01; Electron Microscopy Sciences) were coated with 0.1% (w/v) poly-L-lysine (cat. # P8920; Sigma-Aldrich) in water for 12 hr, then air dried at room temperature.

Sectioning: To allow optimal bonding of the section to the coverslip, the TMA paraffin block face was hydrated in an ice water bath for 3-5 minutes prior to sectioning at 4 µm, then air-dried overnight; well-prepared air-dried sections of BSA gels appear uniformly translucent.

Antigen retrieval: Coverslips were baked overnight at 70°C, deparaffinized in 3 changes of xylene (5 min each), 2x 100% EtOH, (1 min each), 2x 95% EtOH (1 min each), 1x 70% EtOH (1 min) and 2x H_2_O (1 min each). Antigen retrieval was done using Target solution pH 9 (Agilent Technologies) for 20 minutes @ 95°C or 99°C. The solution was preheated to 65°C, coverslips were added, temperature increased with a 20 min ramp to the target temperature, held for 20 min, then cooled to 74°C over ∼ 20 min.

Photochemical bleaching: Bleaching solution was 5% H_2_O_2_, 27 mM NaOH in phosphate-buffered saline (PBS). Coverslips were immersed, cut sections facing up, in transparent shallow dishes sandwiched between two 8.8 x 12.9 cm LED light panels (3.5E4 lux) for 2 cycles of 45 min each, using fresh solution for each cycle (38).

Counterstain: Sections were stained with DAPI (4′,6-diamidino-2-phenylindole) at 0.2 µg/mL in 1X PBS, 20 min, at room temperature.

Antibody staining: After antigen retrieval and photobleaching, the coverslips were mounted sample side up in a metal frame adapted from the Codex/PhenoCycler fluidics system (Akoya Biosciences, Menlo Park, CA) by removing the fluidics tower, and covered immediately with 1 mL antibody diluent buffer: PBS, 3% BSA, 10% goat serum, 0.1% ProClin 300, 0.1% Tween-20, 0.5 mM n-propyl gallate (from 50 mM stock in DMSO; 1% DMSO final concentration). For IHC time course imaging, the antibody concentration was increased in three steps (e.g., 1 nM, 3 nM, 10 nM) over a range intended to bracket the antibody K_D_.

### Imaging

Antibody binding was evaluated at room temperature in real time using a Keyence BZ-X810 inverted microscope with multi-point time-lapse software, motorized stage, metal halide or LED light source each used at 100% power, 20x 0.75 NA plan apochromat objective (Keyence) and an integrated camera; images are 16-bit, 1920 x 1440 pixels at 2.65 pixels/µm. The camera was set to maximum (16x) gain without binning, with dark noise correction off. DAPI and Cy5 filters were Keyence part #OP-87762 and OP-87766, respectively. To minimize non-specific fluorescence from buffer overlying the sample, a custom adapter (Fig. S1) held a 2 mm diameter stainless steel right cylinder with a polished lower end (the “immersion probe”) centered on the optical axis, adjustable on the Z-axis to a position 30-50 µms above the surface of the coverslip. Keyence image metadata includes the Z-axis position of the objective lens. By capturing focused images of both the immersion probe face and the samples mounted on the coverslip, the depth of antibody solution in the field of view can be determined and used to calculate the antibody specific fluorescence (see below).

The imaging sequence (specifying X-Y stage locations, filters and exposures) was defined in the Keyence multi-point time-lapse software. X-Y stage locations were selected to image antigen-containing samples (e.g., BSA gel containing BCL2 peptide, or Granta-519 cell expressing high levels of BCL2) and negative control samples (e.g., BSA gel lacking antigen, or RAMOS cells having minimal BCL2 expression) on the same coverslip. Images of TMA cores were framed to include adjacent glass coverslip areas (“*cis*-glass”) lacking cells or tissue. Additional images were obtained at 4 glass-only reference fields of view located outside the boundary of the TMA cores. To allow image registration at successive time points, DAPI images were obtained at each location at one exposure (typically 2.5E-3 s for cellular samples; 2.5E-2 s for BSA-peptide gel samples). To maximize sensitivity to low-intensity signal while avoiding pixel saturation as the sample fluorescence increased over time, Cy5 images for each location were obtained at 2-3 exposures, typically 0.02-1 s. Between imaging cycles, the stage was positioned so that the immersion probe was located away from the TMA to allow unimpeded antibody access to the TMA cores.

Before beginning antibody incubations, with the sample immersed in antibody diluent buffer, replicate well-focused images were obtained of the immersion probe face, and of samples at each X-Y stage position. The staining time course was then started by removing the diluent buffer, rinsing the TMA with 0.5 mL of the first antibody solution, then adding 1 mL of fresh antibody solution and ∼ 1.2 mL light mineral oil USP to prevent evaporation. For each antibody incubation, images were typically obtained at 30-40 time points at intervals varying from 5 minutes to 6 hours (image acquisition can be less frequent as the reaction proceeds towards equilibrium; details in Table S3). Incubations at each antibody concentration were long enough to allow staining to approach equilibrium, typically 18 to ≥ 48 h.

### Assessment of photobleaching

Photobleaching was assessed using synthetic control sections containing 12.5% BSA and 3E-7 M BSA-AlexaFluor™-647 treated as for experimental samples. Cumulative exposures of 1.5 s per imaging cycle in the Cy5 channel result in photobleaching of ∼1.2% per imaging cycle. DAPI channel exposures up to 0.04 s do not appreciably increase the loss of Cy5 signal. Photobleaching is mitigated by reversible binding of the antibody-dye conjugate, allowing equilibration with unbleached molecules.

### Image registration

Successive images of the same ROI may have translations of varying magnitudes in the X-Y axes. For cellular images, image registration was performed using the MATLAB function imregtform (optimizer parameters: InitialRadius = 0.002; Epsilon = 1.5e-4; GrowthFactor = 1.01; MaximumIterations = 100; metric.NumberOfSpatialSamples = 500). For low-contrast images of uniform BSA-peptide gel samples, the translation data for an adjacent cellular TMA core was used.

### ROI quantification

For BSA-peptide samples, circular ROIs 100 pixels (∼ 38 µms) in diameter were defined. For images of cells, ROIs 70 pixels ( ∼ 26.4 µms) square were defined. Control *cis*-glass ROIs were 70 pixels square or rectangles of similar area. An ImageJ macro opens each image file, selects the desired ROI, adjusts for X-Y translation based on the appropriate image translation data file (control *cis*-glass ROIs were not translated), records the ROI mean fluorescence intensity and the maximum value for any pixel in the ROI. If the maximum pixel intensity is saturated (65535), a software flag is set for that image (SatFlag=0). The resulting ImageJ data is copied into a .csv file for further use by MATLAB software.

### Image analysis

Analysis of the time course data required careful attention to three sources of non-specific signal in the images (39), particularly at early time points when the antigen-specific signal is minimal: 1) variation in blank-field signal arising from non-uniformity in the optical path and from the out-of-focus face of the immersion probe; 2) antibody-containing buffer overlying the sample; 3) antigen-independent binding of antibody to the sample of interest.

### System noise corrections

Under the conditions of these experiments, the integrated camera in our Keyence microscope records a low-intensity (<1% of peak signal), non-uniform, sample-independent “system ghost” signal with exposures of minimal duration and/or samples lacking fluorochrome. To correct for this, each experimental ROI was measured in a “system ghost” reference image averaged from eight 1E-3 s exposures of buffer lacking fluorochrome (Fig. S2), scaled appropriately, then subtracted from the raw intensity value for all other images of that ROI.

### Illumination field heterogeneity

Epifluorescent illumination of a volume of fluorochrome-containing buffer typically creates a radially symmetric, centrally bright image (“vignetting”). The immersion probe presents a polished face perpendicular to the optical axis that produces other nonuniformity due to fine scratches (dark) or contaminating material (variably bright) on the probe face. To account for resulting variation in the intensity of sample illumination and optical path noise, a reference glass ROI image obtained in minimally fluorescent antibody diluent buffer was subtracted from an image of the same ROI obtained in antibody-containing buffer (creating “flat-field image 1”). This step removes the contribution of signal due to illumination of the probe face, which is unchanged in the two images. Remaining variation in image intensity is assumed to arise from variable illumination of the fluorochrome-containing buffer, and to correspondingly affect the signal from a fluorescent sample in the field of view. Accordingly, a normalized flat-field image was calculated by dividing each pixel of flat-field Image 1 by the average pixel intensity of the entire flat-field 1 image, creating a reference flat-field image 2 with pixel values centered around 1 (Fig. S2). For each ROI in all other images, the average value of that ROI in flat-field image 2 was measured to obtain a relative intensity correction factor (RI). For sample ROIs, the raw pixel intensity value, after subtracting the system ghost value, was divided by the RI value before further calculations were made.

### Lamp intensity variability

Variation in sample illumination due to changes in lamp brightness affects the measured signal in every ROI in an image. To account for this, a minimum of three ROIs in “*cis*-glass” areas of buffer solution adjacent to the sample were defined in each image. Because the concentration of fluorescent antibody in the buffer solution is expected to change only gradually during each incubation, short-term image-to-image variation in the measured intensity of these areas is assumed to be due to fluctuation in lamp brightness. Accordingly, for each antibody concentration, ROI image intensities at each timepoint were corrected for the system ghost (SG) and relative intensity (RI) signals, then fitted with a second-order polynomial curve (fluorescence intensity vs. time). Deviations from the polynomial curve at each time point were attributed to lamp intensity variation if they were correlated in the replicate *cis*-glass ROIs in each image; deviations not correlated with other *cis*-glass ROIs in the same image were assumed to be local noise, e.g., from extraneous particles drifting into one ROI. The measured lamp intensity variation at successive time points (typically < 5% from the mean) was used to correct the observed fluorescence intensity values for all other ROIs in the same image.

### Antibody specific fluorescence

To quantify experimental sample fluorescence in terms of the chemical concentration of bound antibody, the specific fluorescence of known concentrations of antibody-fluorochrome in solution was measured using the same optical path, camera and buffers used to measure samples during the actual experiment. In each experiment, the mean fluorescence intensity was measured in ROIs drawn in reference glass-only images of antibody diluent alone, and of each antibody staining solution (typically 1, 3 and 10 nM). The optical path length through the buffer solution in these images was determined from the difference in the z-axis positions of the immersion probe face and the cover glass surface. After correcting for non-specific sources of signal as described above, the mean antibody-specific fluorescence per second of exposure, per µm of path length was calculated using a MATLAB least squares fitting algorithm, assuming a linear correlation between antibody concentration and fluorescence intensity.

Because the coverslip surface is at the focal point of the lens, while the overlying buffer is variably out of focus, the measured fluorescence of the overlying solutions is underestimated relative to the antibody bound to the sample of interest on the coverslip. To account for this, the specific fluorescence of buffer solution for varying optical path lengths was determined by adjusting the gap between the probe face and the coverslip surface. The results show ∼10% underestimation of specific fluorescence at the 30-50 µm buffer optical path length used in these experiments, relative to specific fluorescence in the 5-10 µm path length above the coverslip surface. The specific fluorescence intensity of the antibody was adjusted accordingly.

### Buffer fluorescence and non-specific signal corrections

Fluorescence signal in a sample ROI derives from molecules in both the tissue and the overlying buffer. To correct for the buffer-specific signal, the mean fluorescence intensity of replicate *cis*-glass ROIs was subtracted from the mean fluorescence intensity of sample ROIs. This calculation is done for both antigen-containing and control (no-antigen) sample ROIs. To account for non-specific signal arising from the out-of-focus probe face, the mean fluorescence intensity of each ROI was measured in reference glass-only images, corrected as for the sample ROIs, then subtracted from the sample ROI values. To account for non-specific antibody binding to the sample, the no-antigen control sample fluorescence values were subtracted from the antigen-containing sample values (Fig. S3).

Specifically, for each Ag-containing sample ROI, the raw image intensity was corrected for system ghost noise, optical path non-uniformity, exposure time and lamp intensity as described above. Antigen-independent (non-specific) signal arising from buffer and negative control tissue was calculated combinatorially, according to the number of *cis*-glass and no-antigen control ROIs. For example, a single antigen-containing ROI with 3 adjacent *cis*-glass ROIs, referenced to a no-antigen sample image having 4 negative control ROIs and 3 *cis*-glass ROIs, would yield 3 * 4 * 3 = 36 background-corrected values. (See schematic in Fig. S3) Outliers at each time point (typically due to particles drifting into an ROI) were programmatically excluded using the MATLAB default of >3 times the scaled median absolute deviation (MAD), then the mean +/-2 standard deviations of the remaining unique values was calculated.

The calculation order is as follows:

1. ROIs with saturated pixel values were excluded if the ImageJ SatFlag value = 0; if a shorter exposure of the same ROI was obtained, that image was then examined;
2. ROI values were corrected for system ghost noise and illumination uniformity;
3. ROI values were corrected for exposure time;
4. ROIs were corrected for temporal lamp intensity variation;
5. Signal from *cis*-glass ROIs was subtracted from the sample ROIs in the same image;
6. For all Ag-containing and non-Ag-containing ROIs, the signal from the corresponding reference glass-only ROI was subtracted from the sample ROI signal;
7. Signal from the no-antigen control ROIs was subtracted from the antigen-containing sample ROIs.

At each antibody concentration, a local (moving median) 3*scaled MAD filter (n=7) was used to exclude values affected by transient noise at one time point relative to adjacent timepoints. Excluded data points are plotted on the x axis. Corrected fluorescence intensity was converted to molar concentration of antibody using the previously calculated antibody specific fluorescence. Results are saved to a .csv file (“stats.csv”) along with the details of the ROIs used for the calculations.

### Calculation of limit of blank and limit of detection

The assay limit of blank (LOB) and limit of detection (LOD) were calculated according to established criteria (40). In each of 2 experiments, three negative control cores were compared pair-wise (n=6 comparisons in total), treating one core as “Ag-containing”. For 8-12 ROIs per sample pair (n=50-66 total), at each time point in the first Ab incubation (n=43-49), the absolute value of the mean and the SD of the apparent [AgAb] were determined. An overall mean and SD (mean_global_, SD_global_) of the aggregate data was calculated, taking into account the variability within and between groups. The LOB = mean_global_ + 1.645*SD_global_.

Similarly, in each of 2 experiments, three Ag-containing cores were compared pair-wise with negative control cores (n=6 comparisons in total). The mean and SD of the apparent [AgAb] at each time point in the first Ab incubation (n=43-49) were used to calculate LOD_test_ = mean-1.645*SD. The resulting values were sorted. The LOD was the [AgAb] at the midpoint of the lowest contiguous data set having >95% of its LOD_test_ values greater than the previously identified LOB of the corresponding sample type (BSA gel or cell block).

### Parameter estimation from experimental data

Reaction-diffusion time courses for a range of candidate parameter sets were rapidly predicted by interpolation in a parameter space defined by 2294 pre-calculated reactions. The predicted time courses were compared with experimental data to identify the parameters most consistent with the observed results. See Supporting Material for details and GitHub for MATLAB and ImageJ code.

### Graphs, Statistical tests

All reported results are based on analysis of replicate Ag-containing and negative control ROIs (n=8-12 as detailed in figure legends) in each of replicate TMA cores (n=2-3) in replicate experiments (n=2). Welch’s t-test was performed using GraphPad Prism version 10.5.0 for Windows (GraphPad Software, Boston, Massachusetts USA, www.graphpad.com). Box-and-whisker plots are based on the optimum fitted parameter values obtained for each of 6 candidate [Ag]_total_ and K_D_ values tested for each Ag-containing ROI (n=48-72). Whiskers on box-and-whisker plots show the 10^th^ to 90^th^ percentile range.

### Wasserstein metric

The 2D Wasserstein distance metric quantifies the extent to which an ROI deviates from the assumption of uniform antigen distribution. The metric was minimally adapted from the Wasserstein “multilevelOT” function (41, 42). It takes as input a square matrix of any dimension, with values representing the pixel intensity of the image ROI. The scale of the calculated metric is 0-1, with 0 indicating a completely uniform image intensity, and 1 indicating a maximally non-uniform intensity/spatial distribution, achieved by having a single outlier pixel in a corner of an otherwise uniform image. A single row of positive pixels across the edge of the image yields a value of ∼0.5.

Input data were obtained from 70-pixel square ROIs of Granta-519 cell images taken the end of the staining time course. Non-specific buffer and antigen-independent binding were accounted for by subtracting an image of BSA gel lacking antigen, after first applying the same system ghost, exposure time and relative intensity corrections. ROI pixel values were exported using the ImageJ Image/Transform/Image_to_Results function for analysis in MATLAB.

### Surface plasmon resonance (SPR) analysis

SPR kinetic assays (n=4) were performed using a Biacore T200 (GE Healthcare). Clone 124 mAb (anti-BCL2) was purchased from Abnova (Cat.# MAB1132). The BCL-2 peptide sequence was identical to that used in the BSA-peptide control samples except that the C-terminal cysteine-amide was replaced with a C-terminal lysine(biotinyl)-amide. Peptides were captured on a Biacore Sensor Chip SA (Cat # BR-1005-31; GE Healthcare) in HBS-EP+ (10 mM HEPES, pH 7.4, 150 mM NaCl, 3 mM EDTA, 0.05% Surfactant P20) at 25°C using 1 ng/mL peptide at a flow rate of at 10 µL/min for 60 s. Capture levels ranged from <2-12 RU. Assays were performed at 37°C in 50 mM Tris-HCl, 0.1% BSA (w/v), 0.1% ProClin300 (v/v), pH 7.7. Kinetic analysis was performed using a single-cycle approach. Antibody samples were prepared by 3x serial dilution in running buffer from 60 nM to 0.740 nM, and injected in increasing concentration over a freshly prepared surface. Association was observed for 420 s at 50 µL/min flow; dissociation was observed for 1800 s. Data was fit using Biacore T200 Evaluation software (GE, Version 3.0) with a bivalent analyte model.

### Flow cytometry

Granta-519 and RAMOS cells were obtained from the Genentech cell line repository. Both cell lines were maintained in RPMI 1640 supplemented with 10% FBS (Sigma), pen-strep and 2 mmol/L L-glutamine at 37°C, 5% CO_2_, in a humidified incubator, and harvested at 70-80% confluence. For intra-cellular cell staining, cells were centrifuged, washed and resuspended in PBS at 1E7 cells/mL. Cells were seeded at 1E6 cells per well in a 96-well V-bottom plate, stained with the fixable viability dye eFluor 450 (Invitrogen) for 10 minutes at 4°C, washed, then resuspended in BD fixation/permeabilization buffer (BD Biosciences), incubated for 30 min at 4°C, washed and resuspended in Human TruStain FcX blocking buffer (BioLegend) diluted 1:20 in 1X BD Perm/Wash buffer (BD Biosciences) for 15 min at 4°C. Cells were then resuspended in anti-BCL2 clone 124 AlexaFluor™-647 conjugate, diluted in 1X BD Perm/Wash buffer (BD Biosciences) at a range of concentrations from 1E-9 to 5.4E-8 M, for 30 minutes at 4°C. Cells were then resuspended in 100 µL 1XPerm/Wash buffer and analyzed on the Symphony Flow Cytometer (BD Biosciences).

Cell diameters were measured using a Vi-CELL cell viability analyzer (Beckman Coulter). Volumes were calculated assuming spherical geometry. Fluorescence calibration was achieved using Quantum™ Simply Cellular® anti-Mouse IgG antibody capacity kit (Bangs Laboratories). Quantification beads staining was performed in parallel with the test cell samples. Vendor-provided data was used to establish a correlation between anti-Fc copy number per bead and fluorescence signal; antibody-specific cell fluorescence was interpreted as BCL2 protein copy number by reference to the standard curve. BCL2 concentrations were calculated from protein copy number per cell and cell volumes, assuming that non-specific binding correlates with cell volume.

Representative IHC Keyence image data sets are available for download at Mendeley Data (DOI 10.17632/hvgwj9mn92.1). Representative Virtual Cell models and MATLAB software (both source code and compiled for Windows and Mac/Intel computers) for modeling reaction-diffusion behavior and for image analysis are available at https://github.com/Genentech/IHC-reaction-diffusion.

## RESULTS

### A general model of IHC reaction-diffusion kinetics supports fast interpolation

Our analysis of IHC reaction kinetics proceeded from *in silico* modeling to experimental confirmation. Numerical simulation was first used to model ∼2300 IHC AgAb reactions in uniform samples with different values of [Ag], [Ab], k_for_, k_rev_, x and D_eff_. Each IHC time course was compared to that of a pseudo-first order solution-phase reaction at constant [Ab] with the same [Ag], [Ab], k_for_, and k_rev_. *Iota*_*f*_ was defined as the rate at which the IHC reaction reached a fractional progress, *f*, towards equilibrium relative to the corresponding solution-phase reaction. *Iota*_*f*_ has a value between 1 (*i*.*e*., the IHC reaction proceeds exactly as fast as the solution-phase reaction) and 0 (*i*.*e*., the IHC reaction proceeds infinitely slowly).

Inspection of the data showed that *iota*_*f*_ depends smoothly on three unitless variables: a form of Damköhler number, *kappa* (Eqs. 11, 12); Thiele’s *eta* (Eqs. 13, 14); and [Ab]/K_D_. Any set of IHC reaction parameters maps to one point in a 3-D space defined by these variables, and that point has an associated value of *iota*_*f*_ (Fig. 1). *Kappa* is the primary determinant of IHC reaction behavior when the characteristic diffusion time is long relative to the characteristic reaction time (*i*.*e*., when *kappa* > 1), accounting for the fact that AgAb complex can form only where antibody has diffused into the sample. *Eta* describes the balance between Ab diffusive flux across the sample surface and the AgAb reaction rate in the underlying space; it is the primary determinant of reaction behavior when *kappa* < 1. [Ab]/K_D_ is relevant because antibody diffusion into a sample necessarily creates a transient concentration gradient of free antibody ([Ab_free_]), and therefore a gradient of [Ab_free_]/K_D_, across the depth of the sample. Because the fraction of antigen bound by antibody depends on [Ab_free_]/K_D_ (Eq. 2), the progress of the reaction towards equilibrium depends on the fraction of the tissue volume and reaction duration for which [Ab_free_] is above or below K_D_.

**FIGURE 1.**
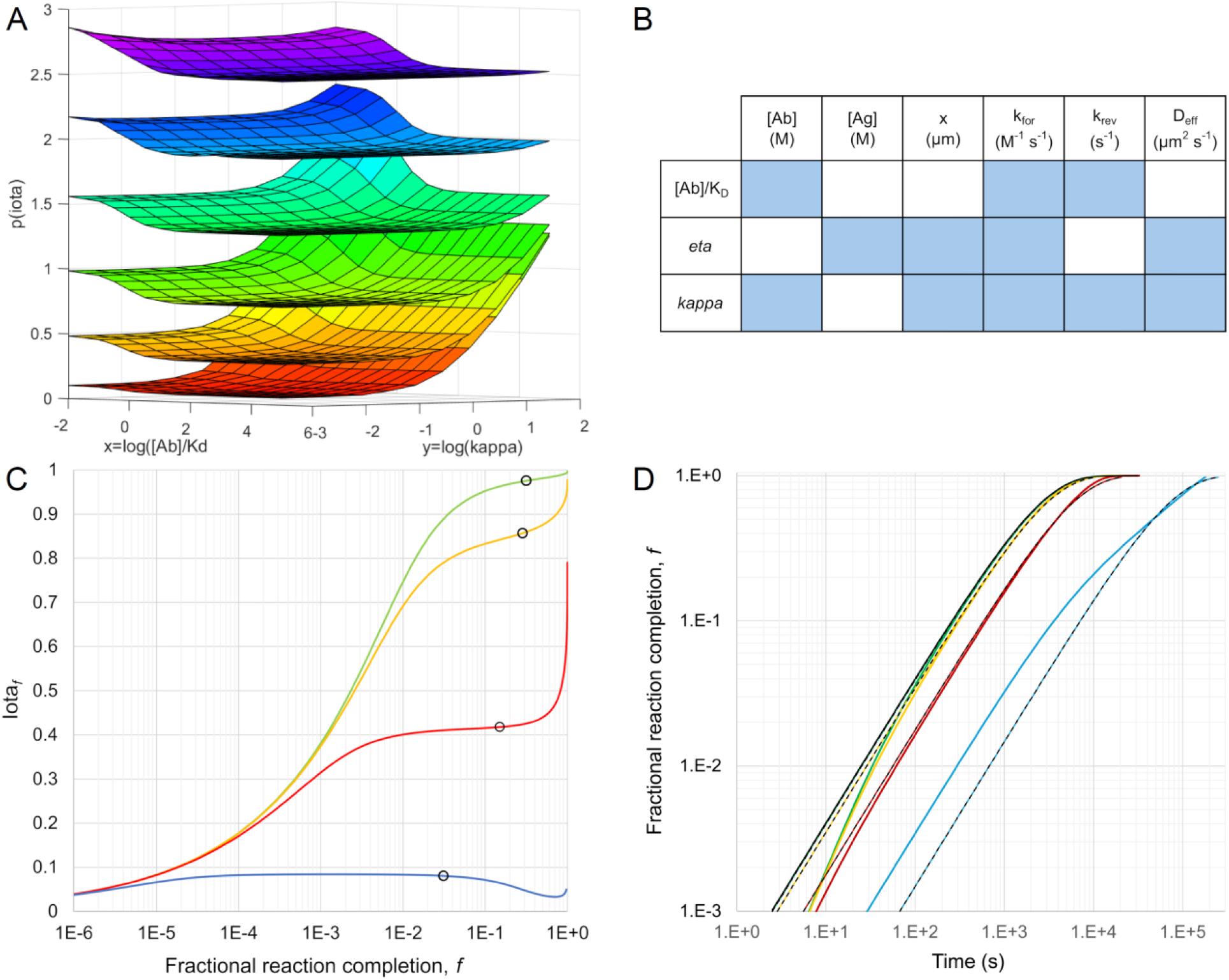
Four unitless parameters characterize AgAb reaction-diffusion kinetics. (*A*) p(*iota*) [-log_10_(*iota*); Z-axis] is plotted as a function of [Ab]/K_D_, *kappa* (X and Y axes, respectively), and p(*eta*) [-log_10_(*eta*); curved leaflets with values of 0.1 (lower leaflet) to 1.6 (upper leaflet), at intervals of 0.3]. Other parameters: [Ab]= 6.67E-9 M (1 μg/mL), [Ag]=2.5E-6 M, k_for_= 1E6 M^-1^s^-1^, k_rev_ = 1E-4 s^-1^, x = 0.5 µm, D_eff_ = 1E-2 µm ^2^s^-1^; *f* =0.8. (*B*) Parameters contributing to [Ab]/K_D_, *eta* and *kappa. Eta* is independent of [Ab]; *kappa* is independent of [Ag]. (*C*) *Iota* varies as IHC reactions proceed towards equilibrium. Antigen concentrations, *eta*, and time to 90% IHC reaction completion are i) 2.5E-8 M, 0.982, 1.59 hr (green); ii) 2.5E-7 M, 0.847, 1.70 hr (ochre); iii) 2.5E-6 M, 0.414, 2.93 hr (red); and iv) 6.25E-5 M, 0.084, 41.2 hr (blue). Circles indicate the 16 minute timepoint for each reaction, highlighting the slower progress of reactions with smaller values of *eta* (higher [Ag]). Other reaction parameters: [Ab] = 4.34E-9 M (0.65 μg/mL); k_for_= 4.5E4 M^-1^s^-1^, k_rev_ = 1.65E-5 s^-1^, K_D_ = 3.67E-10 M, x= 0.5 µm, D_eff_= 1E-2 µm^2^ s^-1^, *kappa*_*90*_ = 3.9E-3. (*D*) Fractional reaction completion (*f*) is plotted vs. time for different [Ag] (parameters and colors as for panel *C*). At constant [Ab], the corresponding solution phase reactions (solid black) reach 90% of equilibrium in 5.7E3 s, superimposing on IHC time courses only when *eta* is near 1 [*e*.*g*., reaction i)]. For reactions ii)-iv) with *eta* < 1, a scaling factor can be applied to k_eff_ (Eq. 3) so that time courses have reaction completion times equal to the IHC reactions at a single value of *f (*here, 50%; color/dashed black). However, these alternatives, which parallel the corresponding solution-phase reaction (solid black), do not accurately describe the IHC time courses at other values of *f*.

A notable result of this analysis is that, for many conditions representative of laboratory IHC reactions, *eta* < 1, and the predicted time required to reach equilibrium correlates with the antigen concentration in the sample when all other conditions are constant. This contrasts with the behavior of first-order reactions at constant [Ab] when diffusion is not rate-limiting (Eqs. 3, 4), for which the relative progress towards equilibrium is independent of [Ag]. Further, the slope and curvature of the IHC time course at different values of *f* varies with *eta*. For this reason, when *eta* <1, a single scaling factor applied to k_eff_ will not accurately describe the IHC reaction time course (Fig. 1D).

The parameter space is analogous to a table of logarithms in that it is time-consuming to calculate in the first instance but allows subsequent calculations to be done accurately and quickly. To predict the time course of an IHC reaction, the reaction parameters are used to determine the corresponding X, Y and Z coordinates in the parameter space (Eqs. 2, 11-14). Interpolation in that space provides the corresponding value for *iota*_*f*_, the rate at which the IHC reaction proceeds relative to the corresponding solution-phase reaction. The corresponding solution-phase reaction time (Eq. 8) divided by *iota*_*f*_ yields the IHC reaction time needed to reach the same *f*. The calculation is repeated for selected values of *f* from 0.001 to 0.9. Implemented in MATLAB, this requires 3-4 seconds for a time course that would otherwise need minutes to days to calculate by *de novo* numerical simulation. Source code and executable software for this function is provided as Fast_Rx_Dif on GitHub.

### Fast interpolation enables objective parameter fitting to model experimental data

Fast interpolation makes possible the time-efficient comparison of experimental time course data with multiple simulated reactions to find the parameters giving the best fit. The analysis requires six input parameters: [Ab], x, [Ag]_total_, D_eff_, k_for_, and k_rev_ (with K_D_ = k_rev_ / k_for_). The first two are controlled by the investigator. The remaining four can be experimentally determined in two steps: [Ag]_total_ and K_D_ are first estimated by fitting experimental data with Langmuir equilibrium theory (Eq. 2); using these values, systematic testing of candidate values for the forward reaction rate constant, k_for_, and the characteristic diffusion time, x^2^/D_eff_, can then be used to find those giving the best fit to the experimental data.

Because the interpolation model training data were generated using MATLAB, the parameter-fitting method was tested against reaction-diffusion data calculated by a mathematically independent method. Accordingly, VCell (35,36) was used to model 27 different IHC conditions distributed across the parameter space. Each VCell time course was then compared to the fast interpolation solutions obtained using 841 candidate pairs of k_for_ and x^2^/D_eff_ and the known K_D_ and [Ag]_total_ values, a calculation requiring ∼20 minutes on a single processor. (Source code and executable versions of the program are provided on GitHub as Standalone_curve_fit). Comparing each predicted time course to the reference VCell data yields a residual fitting error, the average of the fractional difference between the observed and predicted [AgAb] values at each time point. The optimal x^2^/D_eff_ and k_for_ values are those that minimize this error. For the 27 model reactions, the average residual fitting error was 2.2 ± 1.5E-4. The fractional difference between the predicted and “true” x^2^/D_eff_ and k_for_ values (Eq. S17) had a root mean square error = 1.7 ± 1.0E-2 (mean, SD; Table S4), confirming that the fitting process is usefully accurate and precise.

While the optimal x^2^/D_eff_ and k_for_ values are useful measures, a more complete view of the fitting solution is obtained from a contour plot, x = k_for_ ; y = x^2^/D_eff_ ; z = -log_10_(average fractional error between observed and modeled time courses), which defines a point of minimum error surrounded by a roughly hyperbolic region of increasing error (Fig. 2). The vertical axis of the hyperbola delineates reaction-limited conditions for which the value of k_for_ can be determined with relative precision, with a wider range of x^2^/D_eff_ values. Conversely, the horizontal arm delineates diffusion-limited conditions with a well-defined range for x^2^/D_eff_ but a wider range of k_for_ values. The location of the optimal reaction parameters in this space depends primarily on the value of *iota*_*f*_. When *iota*_*f*_ ≈ 1, as occurs when *eta* ≈ 1 and kappa << 1, the optimal parameter pair is in the vertical arm of the hyperbola (Fig. 2A); the IHC process is largely reaction-limited, with kinetics similar to those in solution-phase systems. As *iota*_*f*_ decreases from 1, as when *eta* < 1 and/or kappa > 1, the system moves towards the diffusion-limited domain (Fig. 2B-D). Notably, when the optimal reaction parameters are in the diffusion-limited domain, the corresponding vertical domain of the hyperbola lies to their left, with smaller values of k_for_ and x^2^/D_eff_ (compare Fig 2A and 2D). For this reason, failure to account for the impact of *eta*, or for tissue diffusion rates that are slower than buffer-like conditions, may lead to an underestimation of k_for_ when fitting experimental data. Parameter pairs on a contour boundary define reaction time courses, differing in their slope and intercept, that bracket the experimental time course data (Fig. S4). A boundary at a value of 0.7% average error encompasses the k_for_ and x^2^/D_eff_ parameters used to synthesize the model VCell data, confirming that valid candidate parameters for effective diffusion coefficients and forward reaction rate constants can be objectively extracted from simulated experimental data.

**FIGURE 2.**
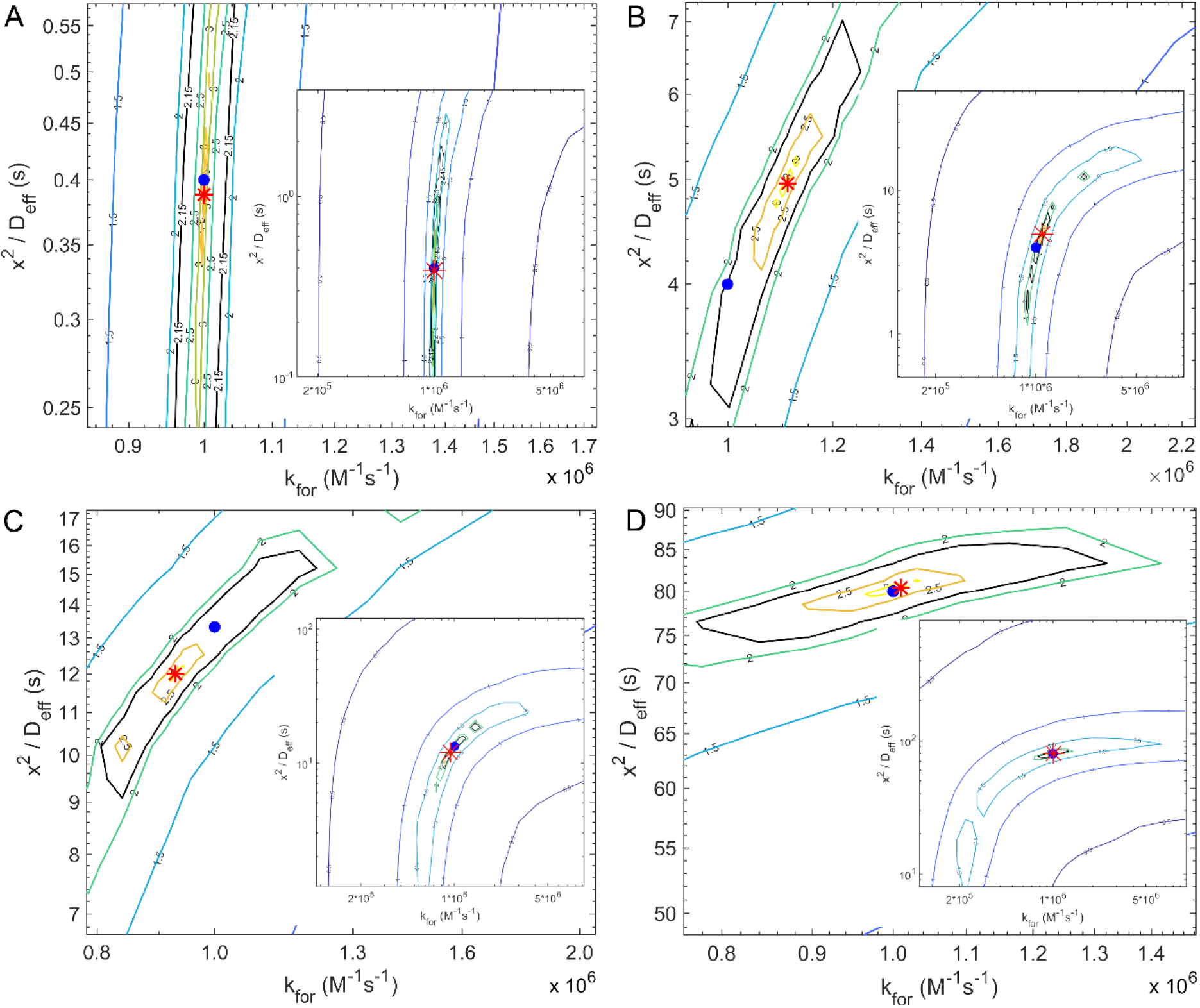
Residual error contour plots define candidate reaction parameter ranges. Contour plots show the negative log_10_ (average fractional difference) between the model VCell time course and the fast interpolation time courses predicted for 841 candidate k_for_ and x^2^/D_eff_ parameter pairs. For all reactions, [Ag] = 1.5E-7 M, [Ab] = 1E-9 M, k_for_ = 1E6 M^-1^s^-1^, k_rev_ = 1E-4 s^-1^, K_D_ = 1E-10 M; x = 4 μm. Blue circles: “true” VCell model k_for_ and x^2^/D_eff_ values; red asterisks: best fit (minimum error) k_for_ and x^2^/D_eff_ values (see Table 2); solid black boundaries at 0.7% average error encompass the true k_for_ and x^2^/D_eff_ values. *A*) D_eff_ = 40 μm^2^s^-1^; the system is reaction-limited; i.e., diffusion provides antibody faster than the antigen binding reaction consumes it; *D*) D_eff_ = 0.2 µm^2^s^-1^; the system is largely diffusion-limited; i.e., the antigen-antibody reaction consumes free Ab as fast as diffusion supplies it.

### Fast interpolation enables objective parameter fitting to synthetic peptide control experimental data

Having confirmed that the model yields appropriate reaction parameters when given simulated IHC data, we next analyzed actual experimental data. The binding of fluorochrome-conjugated anti-BCL2 antibody was measured in 4 µm-thick FFPE sections of well-defined synthetic antigen controls formulated with a 100-fold range of BCL2 peptide (37). A tissue microarray containing cores of these samples was incubated with successively increasing concentrations of antibody (1, 3 and 10 nM), and the change in sample fluorescence was monitored in real time. After appropriately correcting for Ag-independent signal, the resulting data were analyzed to determine [Ag], K_D_, k_for_ and x^2^/D_eff_ as described above.

**TABLE 2.** Actual and fitted reaction parameters for Figure 2.

| Rx | VCell model parameters |  |  |  | Fitted parameters |  |  |  |  |  |
| --- | --- | --- | --- | --- | --- | --- | --- | --- | --- | --- |
| | $D_{\text{eff}}$<br>($\mu\text{m}^2\text{s}^{-1}$) | $x^2 D_{\text{eff}}^{-1}$<br>(s) | $k_{\text{for}}$<br>($\text{M}^{-1}\text{s}^{-1}$) | $x^2 D_{\text{eff}}^{-1}$<br>(s) | $k_{\text{for}}$<br>($\text{M}^{-1}\text{s}^{-1}$) | $k_{\text{rev}}$<br>($\text{s}^{-1}$) | Average<br>fitting error | $\kappa_{90}$ | $\eta$ | $\iota_{90}$ |
| A | 40 | 0.40 | 1.00E+6 | 0.39 | 1.00E+6 | 1.00E-4 | 4.87E-5 | 3.2E-4 | 0.96 | 0.98 |
| B | 4.0 | 4.00 | 1.00E+6 | 4.96 | 1.11E+6 | 1.11E-4 | 5.06E-4 | 3.2E-3 | 0.73 | 0.82 |
| C | 1.2 | 13.3 | 1.00E+6 | 12.0 | 9.28E+5 | 9.28E-5 | 7.22E-4 | 1.1E-2 | 0.48 | 0.62 |
| D | 0.2 | 80.0 | 1.00E+6 | 80.4 | 1.01E+6 | 1.01E-4 | 3.18E-4 | 6.4E-2 | 0.20 | 0.20 |

Analysis of replicate negative control (n=66) and low-positive (n= 50) ROIs from 6 pair-wise TMA core comparisons yielded limit of blank (LOB) and limit of detection (LOD) values of 1.1E-9 M and 1.7E-9 M [Ag], respectively (Table S5). The assay dynamic range between low- and high-positive samples was >1000-fold, with the highest signal corresponding to >2.5 µM [Ag]. Consistent with Langmuir theory (Eq. 2), a stepwise increase in equilibrium [AgAb] was observed with successive increases in [Ab] (Fig. 3), and as predicted by numerical simulations of diffusion-limited reactions, the time needed to reach equilibrium increases with increasing [Ag] (Fig. 3*F*). Under these conditions, qualitative intensity differences in samples with different [Ag] would be observed at all time points, but accurate quantitation of the relative differences in sample [Ag] is possible only when all reactions have approached equilibrium.

**FIGURE 3.**
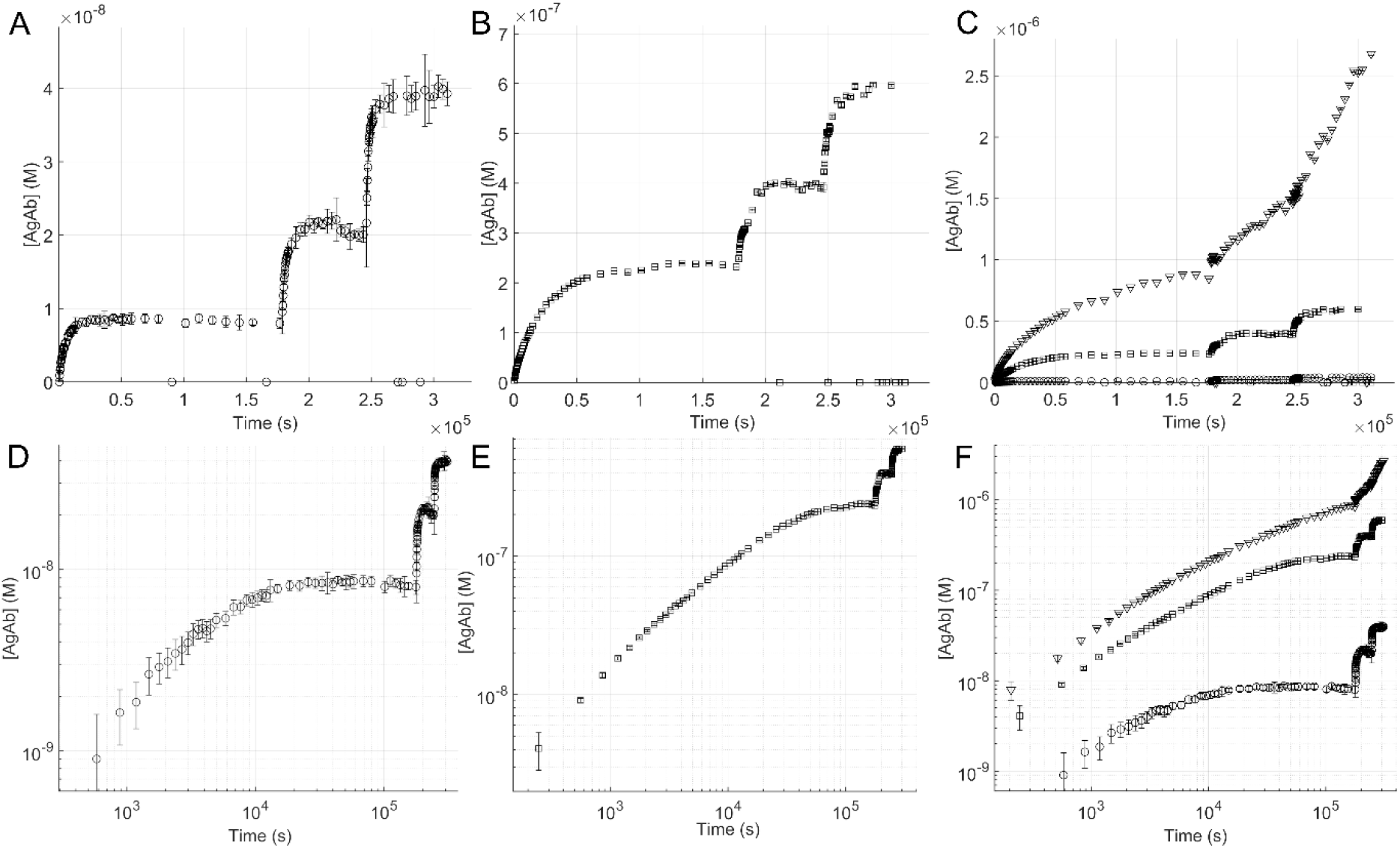
Time to IHC reaction equilibrium correlates with [Ag] in diffusion-limited reactions. Anti-BCL2 Ab binding kinetics measured in synthetic antigen control samples containing BCL2 peptide at 2.5E-6 M (circles; *A,C,D,F*), 2.5E-5 M(squares; *B,C,E,F*) or 2.5E-4M (diamonds; *C, F*). FFPE sections were stained with AlexFluor™-647-conjugated anti-BCL2 clone 124 at successively increasing [Ab]: 1 nM (0-1.8E5 s), 3 nM (1.8-2.4E5 s), and 10 nM (2.4-3.1E5 s) following antigen retrieval at 99°C. (*C, F*) Time course for samples with all three [Ag] are plotted for comparison. Plotted data are the means ± 2 SD of unique values (n=54-144) from combinatorial nonspecific signal corrections. Points on the x-axis (*A, B, C*) are programmatically excluded as outliers relative to adjacent timepoints, or due to saturated pixels (see Methods).

More detailed representative results from one tissue microarray core containing 2.5E-5 M BCL2 peptide are shown in Fig. 4. Quantification of twelve 100 pixel-(38 µm-) diameter regions of interest (ROIs) arrayed on the sample (Fig. 4A-J) yields a K_D_ estimate of 4.1 ± 0.8E-9 M (mean ± SD of 60 unique values estimated for 12 ROIs). For comparison, SPR analysis of clone 124 binding to a peptide containing the same BCL2 epitope (a.a. 41-54) yielded a K_D_ = 8.84 ± 1.62E-9 M (n=4; Fig. S5). Factors contributing to the difference in measured K_D_ may include the physical assay formats, antigen presentation and buffer conditions. The measured average [Ag]_total_ for the 12 ROIs in this experiment was 2.4 ± 0.3E-7 M. The detectable antigen is ∼1% of the formulated BCL2 peptide concentration, consistent with results obtained by us in similar samples using imaging mass spectrometry (37). The difference between formulated and measured [Ag] may be due to chemical modification of antigen by formalin fixation, steric hindrance of Ag-Ab binding or other causes.

**FIGURE 4.**
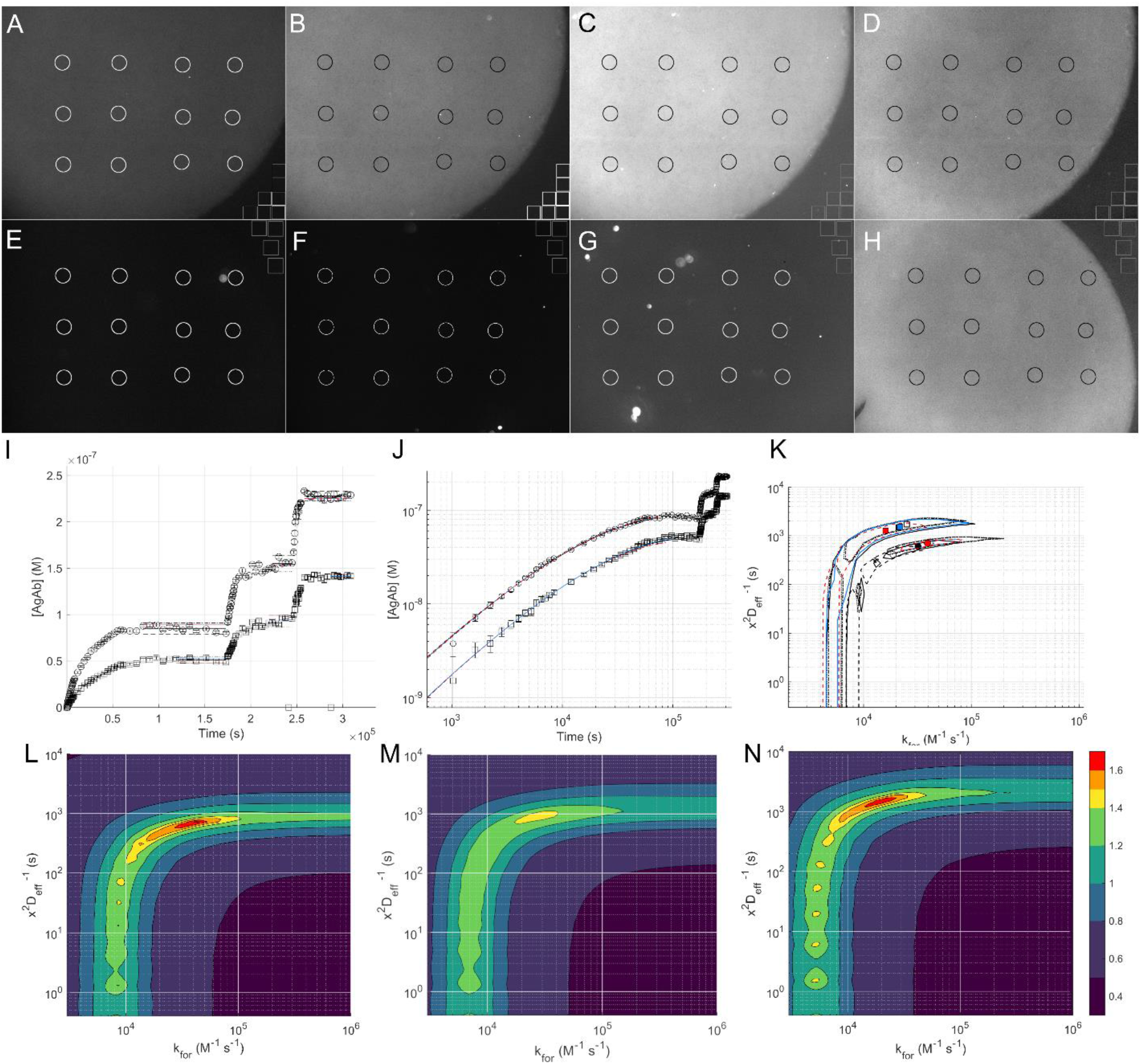
Reaction kinetics can be quantified in well-defined synthetic antigen controls. Shown are representative images of anti-BCL2-AlexaFluor™-647 antibody binding to tissue microarray cores with 2.5E-5 M BCL2 peptide (A-D) and control cores lacking peptide (E-H) after antigen retrieval at 95°C and incubation with three successive [Ab]: (A,E) 9.1E-10 M @ 48 hr; (B, F) 2.5E-9 M @ 68 hr; (C,D,G,H) 9.8E-9 M @ 85 hr. Images A-C, E-G are Cy5 channel, 1 s exposures. Images D and H are DAPI channel (autofluorescence), 0.04 s exposure. All images are shown equally scaled at 0-32768 fluorescent units. Circular regions of interest (ROIs 01-12, top left to lower right) are 100 pixel (38 µm) in diameter. *Cis*-glass control ROIs (n = 7 for Ag-containing cores; n = 4 for control cores) are 70 pixel (26.4 µm) squares. (*I,J*) Observed IHC reaction time courses for ROI 01 (circles) and ROI 12 (squares) are plotted on linear and log scales; data are means ± 2 SD of 420-588 unique background-corrected [AgAb] values at each timepoint; on average, 3.8% of ROI values at each timepoint were programmatically excluded as outliers relative to other values. Horizontal lines (*I*) are [AgAb] plateau values within 2 SD of the mean consistent with: solid black, nominal observed data; red, minimum time course fitting error; blue, minimum Langmuir equilibrium fitting error; black dashed, K_D max_ / [Ag] _max_; dash-dot, K_D min_; dotted, [Ag] _min_. Curves (*J*) are fitted time courses; contour boundaries (*K*) are 3x the minimum fitting error of the optimum parameter pairs (circles, ROI 01; squares, ROI 12). Heat maps (*L, N*) show the average fitting error for the overlapping contour plots (*K*) for ROIs 1 and 12, respectively, and for ROIs 2-11 (*M*). Color map values are -log_10_(average fitting error). Fitting data for ROIs 1-12 are summarized in Table S6.

With these estimates of [Ag] and K_D_, candidate values for k_for_, k_rev_, and x^2^/D_eff_ were determined from the experimental data as described above. The optimal fitted reaction rate parameters (mean ± 1 SD) for the twelve ROIs were: k_for_ : 2.8 ± 1.0E4 M^-1^s^-1^; k_rev_ : 1.1 ± 0.3E-4 s^-1^. For comparison, SPR analysis measured k_for_ = 1.0 ± 0.10E5 M^-1^s^-1^ and k_rev_ = 9.0 ± 2.5E-4 sec^-1^ (n=4; Fig. S5). Parameter fitting yielded optimum x^2^/D_eff_ values of 8.5 ± 3.5E2 s (n = 60; Fig. 4 *K-N*). Fitting errors for the candidate k_for_ and x^2^/D_eff_ parameters are at a minimum in the horizontal diffusion-limited domain (Fig. 4 *K-N*), with x^2^/D_eff_ >5E2 s and k_for_ >1E4 M^-1^ s^-1^. A trend towards slower characteristic diffusion times was noted for ROIs near the edge of the BSA gel core relative to more centrally located ROIs (compare 4N, 4L). The origin of this effect remains for future investigation but may include mechanical compression of the gel matrix during TMA core sampling. With the simplifying assumption that the antibody diffusion distance is the intended section thickness (4 µm), the mean ± 1 SD of fitted x^2^/D_eff_ values (5.0–12E2 s) correspond to D_eff_ values of 1.3-3.2E-2 µm^2^ s^-1^, ∼1000-3000 times slower than the 40-44 µm^2^ s^-1^ reported for IgG in aqueous buffer (25,26).

The results shown above are representative of a larger data set. As expected, analyses of images from paired antigen-containing and negative control (“NoAg”) cores showed similar results (Welch’s t-test; p>0.05) regardless of the choice of NoAg core. Within one experiment, replicate Ag-containing cores yielded parameter values with overlapping ranges whose means differed by up to 40% (Fig. 5). Compared with the experiment shown in Fig. 4, done with antigen retrieval at 95°C, a replicate experiment identical in design except for having antigen retrieval at 99°C shows 2- to 3-fold higher [Ag] values (p < 0.001; Welch’s t-test; Fig 5A). In contrast, the optimum fitted x_2_/D_eff_ and K_D_ values show smaller differences of variable significance between experiments (Fig. 5B, C).

**FIGURE 5.**
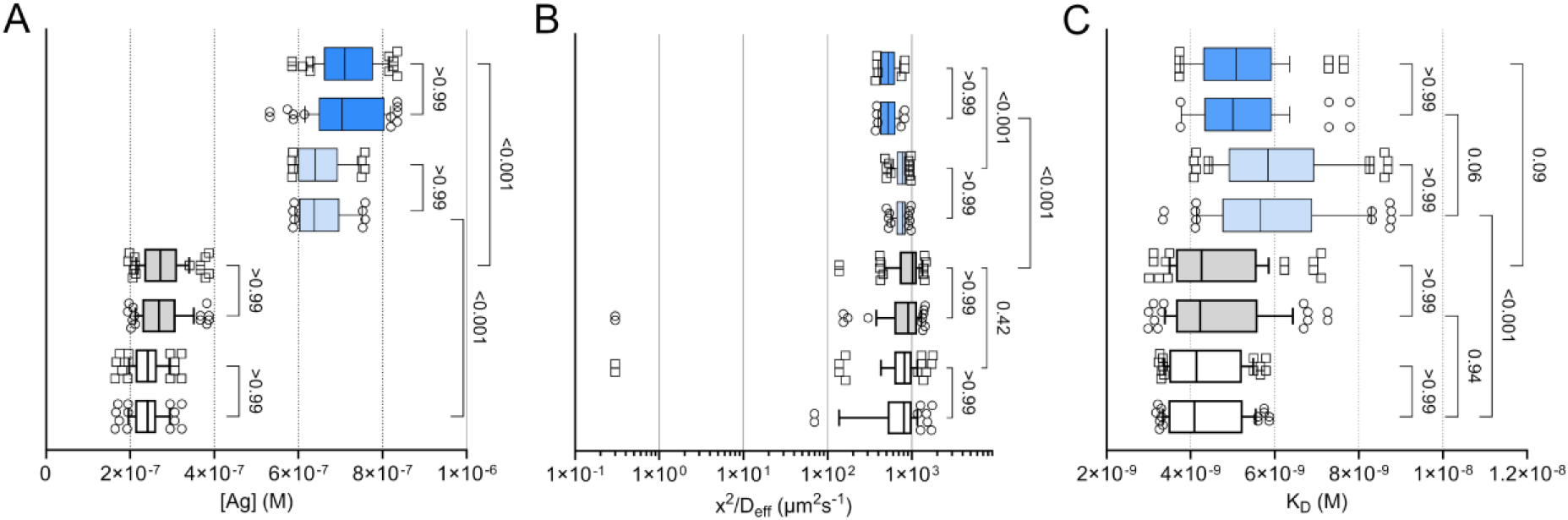
Summary of synthetic antigen control BSA gel analyses. (*A-C*) Data are optimal parameter fits from two experiments, identical in design except for having antigen retrieval at 95°C (white/grey) or 99°C (blue). Whisker plots show 10^th^-90^th^ percentile. For each experiment, results are shown from 4 calculations using replicate “Ag” (white/grey, light/dark blue) and “NoAg” (circle and square outlier data points) images. Lower row (white/circles) are results from Fig. 4. For each data set, n= 48-72 (8-12 Ag ROIs, each with 6 parameter fits).

### Fast interpolation enables objective parameter fitting to cell block experimental data

We next asked if reaction parameters could be measured in fixed cells by examining FFPE sections of BCL2-expressing Granta-519 cells, using RAMOS cells, which lack measurable BCL2 expression, as a negative control sample. Analysis of replicate negative control (n=50) and low-positive ROIs (n = 50) from 6 pair-wise comparisons yielded limit of blank (LOB) and limit of detection (LOD) values of 1.5E-9 M and 2.4E-9 M [Ag], respectively (Table S5). In a representative analysis, eight ROIs measuring 70 x 70 pixels (∼26.4 x 26.4 µm) were quantified and fit to candidate reaction parameters as before (Fig. 6; Tables 3, S7). Background-corrected average fluorescence intensity measurements yield an average BCL2 concentration of 9.4 ± 1.7E-8 M (mean ± SD of 78 unique parameter fitting results for 8 Granta-519 cell ROIs in one TMA core referenced to 2 RAMOS cell negative control cores; light blue box-and-whisker plots in Fig. 7). For comparison, FACS analysis of fixed, permeabilized Granta-519 cells (corrected for non-specific antibody binding to RAMOS cells) yielded 1.6-1.8E5 anti-BCL2 antibodies bound per cell, corresponding to 1.9-2.1E-7 M, approximately 2-fold higher than we measure in FFPE tissue sections (Fig. S6).

**TABLE 3.** Representative fitted parameters for Granta-519 cells.

| Fit | Line | [AgAb]<sub>1</sub><br>(M) | [AgAb]<sub>2</sub><br>(M) | [AgAb]<sub>3</sub><br>(M) | K<sub>D</sub> mean<br>(M) | [Ag]<sub>total</sub><br>(M) | K<sub>D</sub> error | χ<sup>2</sup>/D<sub>eff</sub><sup>-1</sup><br>(s) | k<sub>for</sub><br>(M<sup>-1</sup>s<sup>-1</sup>) | k<sub>rev</sub><br>(s<sup>-1</sup>) | minimum<br>fit error |
| --- | --- | --- | --- | --- | --- | --- | --- | --- | --- | --- | --- |
| nominal | ————— | 4.13E-8 | 6.80E-8 | 9.70E-8 | 4.09E-9 | 1.15E-7 | 7.33E-3 | 1.21E+3 | 1.03E+5 | 4.23E-4 | 1.81E-2 |
| K<sub>D</sub> min | ----- | 4.18E-8 | 7.27E-8 | 8.77E-8 | 2.86E-9 | 1.00E-7 | 2.08E-3 | 1.22E+3 | 1.46E+5 | 4.17E-4 | 2.41E-2 |
| K<sub>D</sub> max | - - - - - | 4.09E-8 | 6.32E-8 | 1.06E-7 | 6.07E-9 | 1.38E-7 | 1.58E-2 | 1.24E+3 | 7.15E+4 | 4.34E-4 | 1.55E-2 |
| [Ag]<sub>min</sub> | ***** | 4.18E-8 | 6.98E-8 | 8.77E-8 | 2.96E-9 | 1.00E-7 | 1.22E-3 | 1.24E+3 | 1.42E+5 | 4.19E-4 | 2.38E-2 |
| [Ag]<sub>max</sub> | - - - - - | 4.09E-8 | 6.32E-8 | 1.06E-7 | 6.07E-9 | 1.38E-7 | 1.58E-2 | 1.24E+3 | 7.15E+4 | 4.34E-4 | 1.55E-2 |
| K<sub>D</sub> err_min | ————— | 4.10E-8 | 7.04E-8 | 8.77E-8 | 3.04E-9 | 1.01E-7 | 2.93E-6 | 1.15E+3 | 1.23E+5 | 3.73E-4 | 1.82E-2 |
| Mean |  | 4.13E-8 | 6.88E-8 | 9.33E-8 | 3.81E-9 | 1.11E-7 | 5.30E-3 | 1.21E+3 | 1.17E+5 | 4.13E-4 | 2.00E-2 |
| SD |  | 4.30E-10 | 3.57E-9 | 8.39E-9 | 1.36E-9 | 1.66E-8 | 6.53E-3 | 3.53E+1 | 3.05E+4 | 2.32E-5 | 3.81E-3 |

**FIGURE 6.**
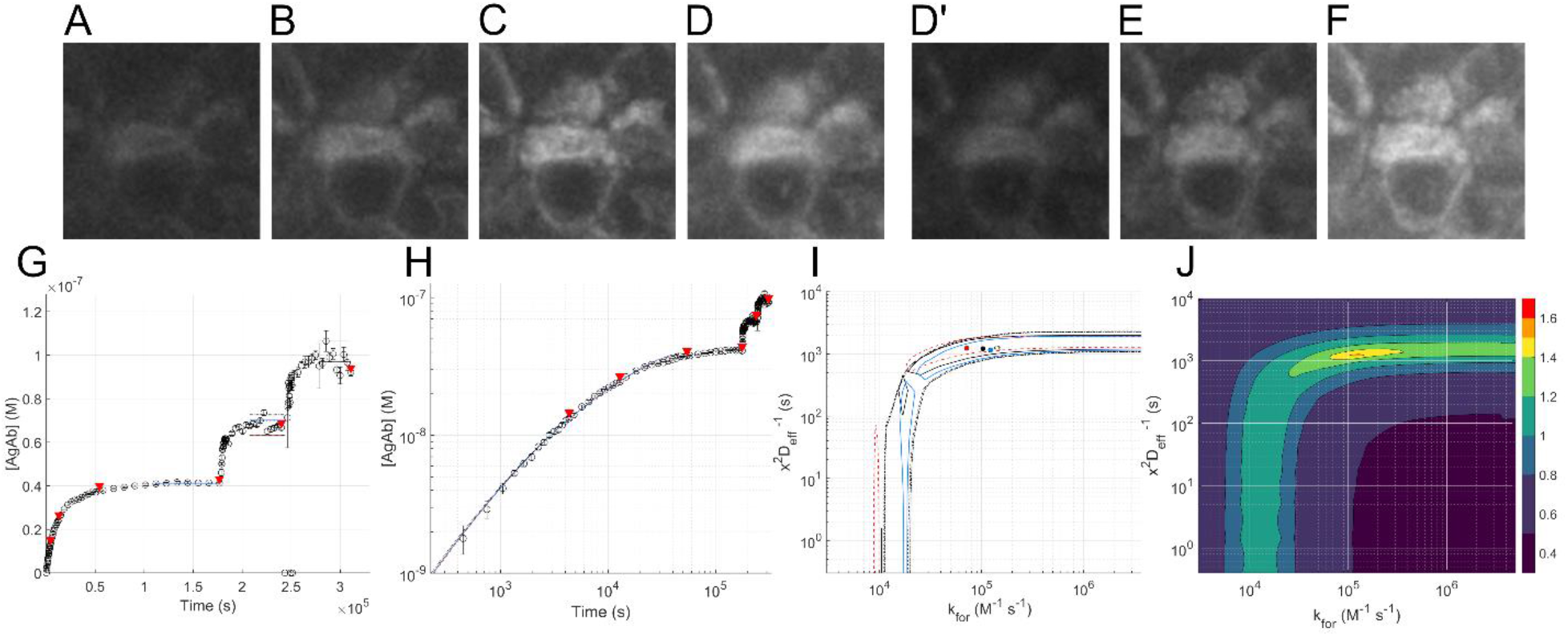
Analysis of Granta-519 cell staining. Representative images are shown of one 70 pixel (26.4 µm) -square ROI after antigen retrieval at 99°C and successive incubation with [Ab] = 1.0E-9 M (*A-D*), 3.5E-9 M (*E*), and 1E-8 M (*F*) at timepoints from 4E3 (*A*) to 3E5 s (*F*) marked with triangles in (*G,H*); image exposures are 1 s (*A-D*) and 0.5 s (*D’-F*); displayed pixel intensity ranges are equal for all images. (*G,H*) AgAb complex formation vs. time is plotted on linear and log scales; error bars are 2 standard deviations for replicate calculations of non-specific signal correction (n = 77-128; mean = 123); points on the X-axis in (*G*) are programmatically excluded outliers relative to adjacent timepoints. (*I*) Contour plot (k_for_ vs. x^2^/D_eff_ ) boundaries at 3 times the minimum fitting error define the range of reaction parameters consistent with the observed time course data; patterned boundaries correspond to the optimum fits defined by parameter combinations as noted in Table 3; circles represent optimal fitted k_for_ vs. x^2^/D_eff_ pairs; black: “nominal”; red: minimum time course fitting error; blue: “K_D error_min_”; open circles, other. (*J*) “Heat map” contour plot shows the average of the overlapping contour plots in I; colormap shows the -log_10_(average fractional fitting error).

**FIGURE 7.**
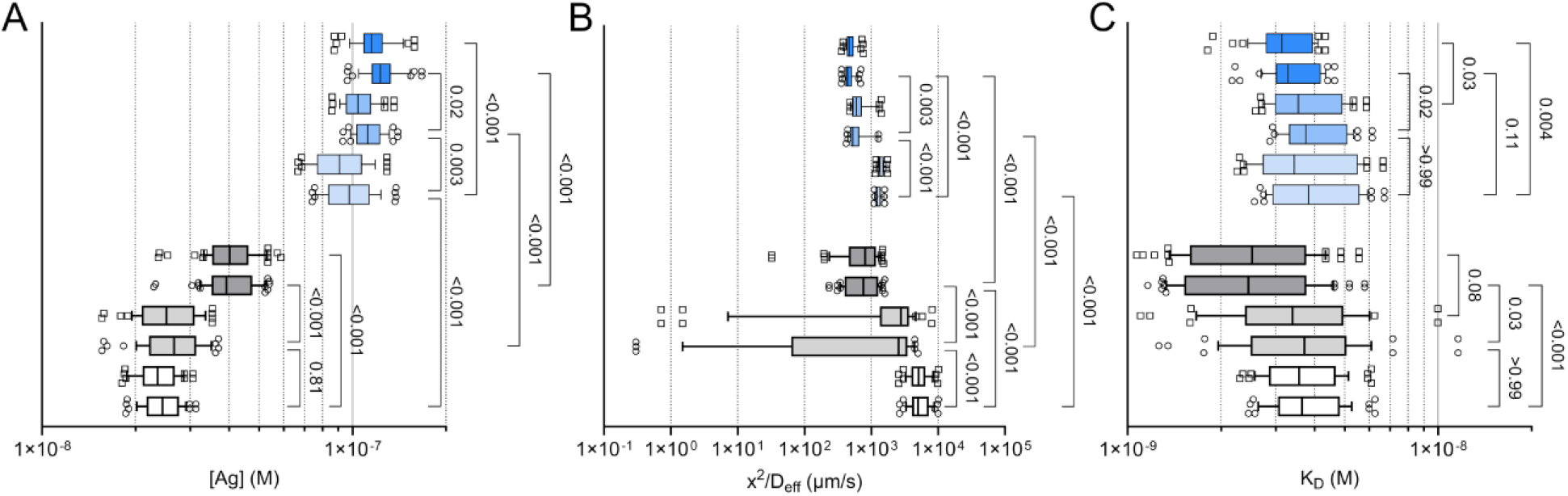
Summary of Granta-519 cell image analyses. (*A-C*) Data are optimal parameter fits from two experiments, identical in design except for having antigen retrieval at 95°C (white/grey) or 99°C (blue). Whisker plots show 10^th^-90^th^ percentile. For each experiment, results are shown from 6 calculations using 3 replicate “Ag” (white/light/dark grey, light/medium/dark blue) and 2 “NoAg” (circle and square outlier data points) images, with 8-12 Ag-containing ROIs per image. For each data set, n= 48-72. Results from Fig. 6 and Table 3 are included as one of the 8 ROIs in row 6 (light blue, circles).

For the 8 ROIs in this Granta-519 core, the measured on-slide K_D_ for anti-BCL2 binding was 3.7 ± 1.2E-9 M (n= 78), similar to the value measured in the BSA-BCL2 peptide synthetic control samples. The mean ± 1 SD of the 78 unique optimum fitted values for k_for_ and k_rev_ were 1.3 ± 0.5E5 M^-1^sec^-1^ and 4.6 ± 1.7E-4 sec^-1^, respectively. The corresponding x^2^/D_eff_ values are 1.3 ± 0.16E3 sec, consistent with diffusion-limited reaction conditions. Given the section thickness (x=4 µm), the corresponding D_eff_ values, ∼1.2E-2 μm^2^s^-1^, are ∼3000-fold slower than reported for IgG (40-44 μm^2^sec^-1^) in buffer (25,26).

As was true for the synthetic antigen control samples, when 2 different “NoAg” cores (RAMOS cell samples) were used as controls for the same Ag-containing Granta cell core, 15 of 18 parameter fitting calculations yielded a similar range of values for [Ag], x^2^/D_eff_, and k_for_ (p > 5E-2); for 3 other cases, the paired results varied by less than 10% (p = 1E-3 to 4E-2). Within any one TMA, the mean observed [Ag] in replicate cores of Granta cells differed by less than 1.6-fold. Notably, similar to what we observed in the BSA-antigen gels, measured [Ag] was consistently 3-to 4-fold higher (p < 1E-3) in cores subject to antigen retrieval at 99°C vs. 95°C (Fig. 7A). In cores subject to antigen retrieval at 95°C, the observed characteristic diffusion times (x^2^/D_eff_) varied 1.4- to 7-fold, a result that may reflect local variation in sample fixation or other preanalytic variables. In contrast, in samples subject to antigen retrieval at 99°C, the range of measured x^2^/D_eff_ values was narrower and the mean values for paired cores were 2- to 4-fold lower (p < 1E-3) (Fig. 7B). The average observed K_D_ values were similar in cores subject to antigen retrieval at 99°C vs. 95°C, varying from 2 to 4E-9 M (Fig 7C). These results support the conclusion that antigen retrieval at higher temperatures may improve Ag detection by increasing both the fraction of Ag accessible to antibody, and the rate at which antibody can diffuse into the sample, without substantially affecting the Ag-Ab K_D_.

### Antigen spatial heterogeneity has limited impact on parameter fitting

Unlike the idealized reaction-diffusion model and the BSA-peptide synthetic control samples, antigen distribution in Granta cells and other biological samples is spatially nonuniform. Modeling shows that diffusion of free antibody in areas with high antigen concentrations may be slower than in areas with lower antigen concentrations, leading high-antigen areas to recruit free antibody from adjacent low-antigen areas, accelerating and slowing the reactions in the respective regions (Figs. 8, S8). To understand the magnitude of this effect in the Granta-519 sample ROIs, we quantified the spatial heterogeneity of antigen distribution in the square ROIs using an adaptation of a 2D Wasserstein distance (WD) metric (41, 42). A WD value of 0 indicates spatially uniform signal; a value of 1 indicates the signal is maximally non-uniform (1 pixel in one corner of the image is positive). Quantification of 50 representative 70 pixel (26.4 µm) -square ROIs of BCL2 staining in Granta-519 cells yielded WD =2.15 ± 0.63E-1.

**FIGURE 8.**
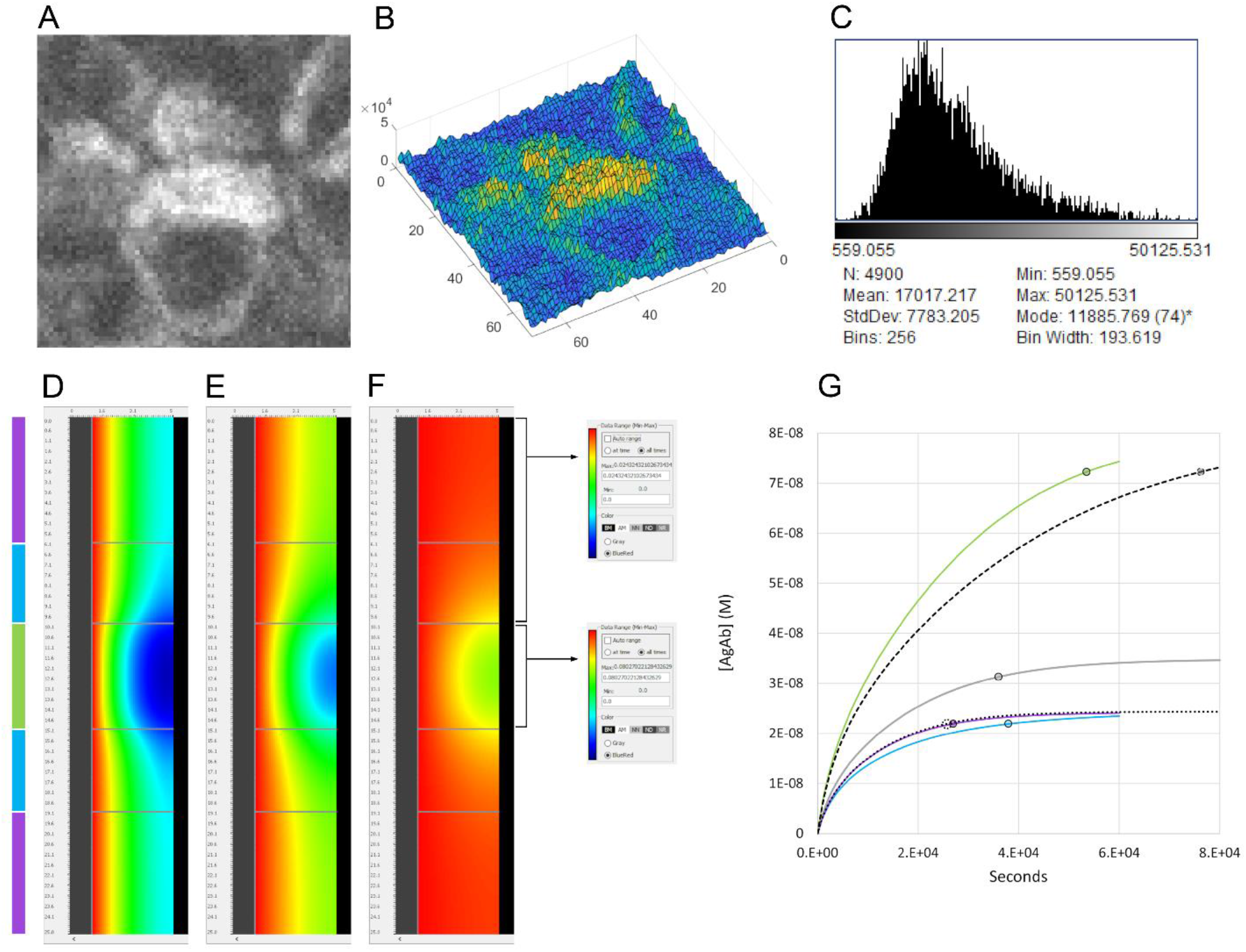
(*A*) Granta-519 cell sample (from Fig. 6) following 48 hours incubation with 1E-9 M Ab, 70 x 70 pixels (26.4 um square), corrected for non-specific signal; (*B*) Surface plot of ROI 104; the scale-invariant Wasserstein distance metric for this ROI is 0.258; (*C*) Histogram of ROI; (*D-G*) VCell model of a 4 µm-deep, 25 µm-wide tissue model with [Ag] = 2.475E-7 M (“high-[Ag]”) in the central 5 µm-wide zone, [Ag] = 7.5E-8 M (“low-[Ag]”) in the lateral 10 µm-wide zones. The Wasserstein distance metric for this geometry is 0.250, at the 78^th^ percentile of 50 ROIs measured in 2 experiments; (*D-F*) distribution of AgAb product at intermediate timepoints illustrates the impact of adjacent high-and low-[Ag] regions on each other; (*D*), 9E3 seconds; (*E*), 1.8E4 seconds; (*F*), 4E4 seconds; see Fig. S8 for movie; (*G*) reaction time courses for different section of the VCell model are plotted; in the high-[Ag] ROI (middle panel in *D-F*, green curve in *G*) the contribution of free antibody from the adjacent low-[Ag] ROI accelerates the reaction progress towards equilibrium relative to a sample with uniform high-[Ag] (black dash); because of this Ab “steal” effect, the adjacent low-[Ag] ROI (cyan) proceeds to equilibrium at a slower rate than the outer low-[Ag] ROI with the same [Ag] (teal). For comparison, time courses for uniformly low and global average antigen concentrations are shown (dotted and grey curves, respectively). Circles indicate times to 90% reaction equilibrium. Other Virtual Cell model parameters: [Ab] = 1E-9 M; k_for_ = 1.2E5 M^-1^ s^-1^; k_rev_ = 5E-4 s^-1^; x = 4 µm; D_eff_ = 1.2E-2 µm^2^ s^-1^.

To understand how the fitting program functions with spatially heterogeneous samples, we created a data set *in silico* using a 2-dimensional Virtual Cell model with antigen spatial heterogeneity (Wasserstein metric = 2.50E-1), average antigen concentrations, k_for_, k_rev_ and D_eff_ values (1.07E-7 M, 1.2E5 M^-1^s^-1^, 5E-4 s^-1^ and 1.2E-2 µm^2^s^-1^, respectively) representative of the results obtained from fitting Granta-519 cell samples. The resulting synthetic time course data were then fit using the global average antigen concentration in the entire 25 µm-wide model as an input reaction parameter. The fitted optimal x^2^/D_eff_ and k_for_ parameters were 112% and 108%, respectively of the actual VCell parameters; the parameter-fitting error contour boundary includes both values (Fig. S7). For comparison, we modeled the system using a second average [Ag] calculated for the 5 µm-wide central core together with the two adjacent low-[Ag] 4 µm-wide ROIs. For this condition, the predicted optimal x^2^/D_eff_ and k_for_ parameters were marginally closer, 108% and 104%, respectively, to the actual Virtual Cell parameters. We conclude that the extent of antigen spatial heterogeneity found in these samples has a limited and quantifiable impact on the parameter fitting results when the average concentration of antigen in the sample ROI is used as an input value.

## DISCUSSION

We demonstrate here the use of practical software planning and analysis tools and experimental methods to quantify the biochemical antigen concentration and reaction parameters determining IHC reaction kinetics in a tissue section. In 4 µm FFPE sections, experimental results for a widely used mouse monoclonal antibody, anti-human BCL2 clone 124, show that the IHC process is functionally diffusion-limited, with the effective diffusion coefficient 1000-3000 times slower than in aqueous buffer.

The observation that IHC reactions may be diffusion-limited has important implications for the interpretation of results: as the [Ag] in a sample increases, more time is needed for diffusion to deliver antibody sufficient to reach equilibrium with the available antigen. If the staining in two samples with different [Ag] is compared before the slower reaction (*i*.*e*., the sample with the higher [Ag]) has reached equilibrium, the observed difference in staining intensity will underestimate the true difference in [Ag], a result we term “dynamic range compression.” (Fig. S9.) To help understand this risk, and to enable more fully informed experimental planning and interpretation, executable software programs are provided to allow users to rapidly model IHC reactions for any set of parameters, and to analyze experimental IHC time course data to objectively find the reaction parameters most consistent with the observed results.

Comparison of results from different staining protocols requires complete documentation of conditions. Per Fick’s first law, the rate of antibody entry into the specimen (formally, “flux”, *J*(*t*) = −*D*_*eff*_ ∗ *d*[*Ab*]_*x*=0_ / *dx* ) is proportional to the effective diffusion coefficient and the concentration gradient between the staining solution and the depth of sample. For any specimen, the higher the antibody concentration in the staining solution, the faster the reaction will reach equilibrium. It is therefore essential to know the molar concentration of antibody applied to the sample. Alternative descriptions including dilution from stock are not informative, and experimental results reported without this value cannot be rigorously interpreted.

Current theoretical descriptions of antibody diffusion and binding are most advanced and mathematically rigorous in the mass transport–limited binding kinetics regime, largely due to decades of work with biosensor systems such as SPR and biolayer interferometry (BLI). These platforms are well-defined (planar surfaces, controlled flow, and simple geometries), making them ideal testbeds for developing and validating models. Our analysis of IHC reaction kinetics shares some common ground with this well-established theory. In particular, *eta* (32) and the mathematically identical “thickness factor”(33) both quantify, under hypothetical quasi steady-state conditions, the impact of diffusion on the forward reaction rate averaged over the entire sample volume, relative to the reaction rate at the surface in the sample.

In other details, however, differences between IHC and SPR reaction geometries and experimental conditions justify different approaches to the analysis of reaction kinetics in these systems. In SPR experiments, antigen is typically bound in a hydrogel surface layer up to 100-200 nm thick with a diffusion coefficient estimated to be 10 to 40-fold slower than aqueous buffer (43). Antibody delivery is controlled by the device channel dimensions, buffer flow rate and antibody diffusion coefficient. Experimental conditions (hydrogel layer thickness, target density, ligand bulk delivery rate) can be adjusted to avoid reagent transport limitations that complicate interpretation of the kinetic parameters obtained (31, 33). In contrast, a 4-5 µm IHC section is ∼20-50 times deeper than the SPR surface layer, with an effective diffusion coefficient 1000 to 3000-fold lower than aqueous buffer (this work). Antigen concentration, determined by the biology of the system, is outside the control of the investigator; section thickness and the effective diffusion coefficient of antibody in the tissue can be modified only within limits determined by the physical integrity of the sample. The approach described here accounts for these IHC-specific conditions, allowing accurate modeling of reaction kinetics across 90% of the reaction time course.

A number of simplifying assumptions should be understood when interpreting the model’s predictions: 1) Tissue antigen concentration is assumed to be spatially uniform, whereas in biological samples it is not. In practice, this has only limited impact; quantification of BCL2 antigen spatial heterogeneity in Granta-519 cells and *in silico* modeling show that the resulting errors in experimental x^2^/D_eff_ and k_for_ estimation are <15% and < 10%, respectively. 2) Rather than being uniform, the diffusion rate and path length for an antibody in a single region of interest may vary due to the presence of large penetrating voids (e.g., lung alveolar air spaces) or smaller channels (e.g., vascular or lymphatic spaces, exocrine ducts). For these reasons, we model the average characteristic diffusion time, combining distance and diffusion variables as x^2^/D_eff_. We expect that the value of x^2^/D_eff_ will further vary according to tissue type, fixation, antigen retrieval method, buffer composition and variables specific to the antibody *per se*. 3) The antibody concentration in the staining solution is assumed to be uniform in space and time, whereas in an unstirred staining solution, a gradient of [Ab] would develop at the tissue boundary (1,31). In practice, our apparatus periodically stirs the overlying antibody solution as the stage moves to different locations, reducing the gradient. Further, while the concentration of free antibody in solution falls during the reaction, the 1 mL staining solution buffer we use typically contains a sufficient excess of antibody relative to tissue antigen that this has minimum impact. 4) While antibodies are modeled as kinetically bivalent (*i*.*e*., with 2 functional Fab domains), we assume they are functionally monovalent (*i*.*e*., only one antigen would be bound per antibody). This assumption is based on the fact that biologically relevant antigen concentrations (typically < 1E-6M), if uniformly distributed in space, result in intermolecular distances greater than the 12-15 nm span between the Fab domains on a single IgG. Formalin fixation cross-links proteins and paraffin embedding removes physiological lipids, preventing lateral diffusion and “patching” of monovalent antigens. However, multivalent antigens, or monovalent antigens cross-linked during sample preparation could nevertheless bind antibodies bivalently. In this event, the observed reaction rate will be faster than the model predicts (44), the observed equilibrium [AgAb] will be reduced by up to 2-fold, and dissociation will be slower than the expected monovalent off-rate. Under these circumstances, more accurate quantification of reaction kinetics and antigen concentration could be achieved with a monovalent antibody reagent (*e*.*g*., Fab, nanobody or similar). 5) Variation in section thickness and dimensional changes incurred during tissue processing (45) will affect the apparent concentration of antigen. 6) Pre-analytic variables including tissue fixation and sample preparation steps, including antigen retrieval, will affect the fraction of antigen that is biochemically intact and accessible to antibodies; these variables should be controlled as much as possible (45,46). 7) Finally, the analysis described here is only one part of quantifying more complex IHC protocols often used in experimental and diagnostic assays. These involve unconjugated primary antibodies detected by incubation with fluorochrome-or hapten-conjugated secondary antibodies, each with their own set of reaction-diffusion parameters. In the latter case, enzymatic chromogen formation further complicates the correlation with [AgAb]. Quantification of these steps would be a valuable accomplishment but is outside the scope of this work.

Despite these caveats, the ability to objectively fit candidate AgAb reaction parameters to a wide range of experimental data - including conditions between the reaction-limited and diffusion-limited extremes, a domain that is intractable to solution using calculus - is a significant advance. False precision is avoided by defining boundaries around the range of parameter combinations most consistent with the observed experimental data, rather than by identifying a single optimum. The approach presented here advances the goal of replacing IHC endpoints that are semi-quantitative and protocol-specific with universally quantitative ones, offering the possibility to correlate results from IHC assays developed with different reagents and protocols at different sites. Potential applications include biomarker discovery, diagnostics, and therapeutic evaluation, where precise quantification and comparison of antigen levels in patient samples may be crucial for meaningful insights and decision-making. More broadly, the principles relevant to IHC analysis apply equally to other tissue-based “spatial omics” analyses, and they should be considered when designing and interpreting experiments requiring any macromolecule to diffuse into and react in a tissue section.

## DATA AVAILABILITY

Representative executable MATLAB code, ImageJ macros and Virtual Cell models are available at https://github.com/Genentech/IHC-reaction-diffusion. Representative Keyence image data are availabe at Mendeley Data (DOI 10.17632/hvgwj9mn92.1).

## AUTHOR CONTRIBUTIONS

W.P., D.D. and F.P. performed and analyzed IHC experiments. B.A. designed, performed and anayzed SPR experiments. X.W. contributed to the design and analysis of SPR experiments. B.M. designed, performed and analyzed FACS experiments. J.E. and H.N. provided software support. A.C. and D.O. advised on IHC experimental design and theoretical analysis, and helped write the manuscript. F.P. wrote modeling and analysis programs, and wrote the manuscript.

## ACKNOWLEDGMENTS

The authors are grateful for the expert advice and technical support provided by many colleagues including Miriam Baca, Cecile Chalouni, Magnus Fontes, Sarah Gierke, Charles Havnar, Gabe Jackman, Hartmut Koeppen, Praveen Krishnamurthy, Serena Y. Lee, Kathryn Mesh, Tatiana Novitskaya, David Peale, John Quinn, Linda Rangell, Dick Vandlen and Andries Zijlstra.

The Virtual Cell is supported by NIH Grant R24 GM137787 from the National Institute for General Medical Sciences.

## SUPPLEMENTAL FIGURES

**FIGURE S1.**
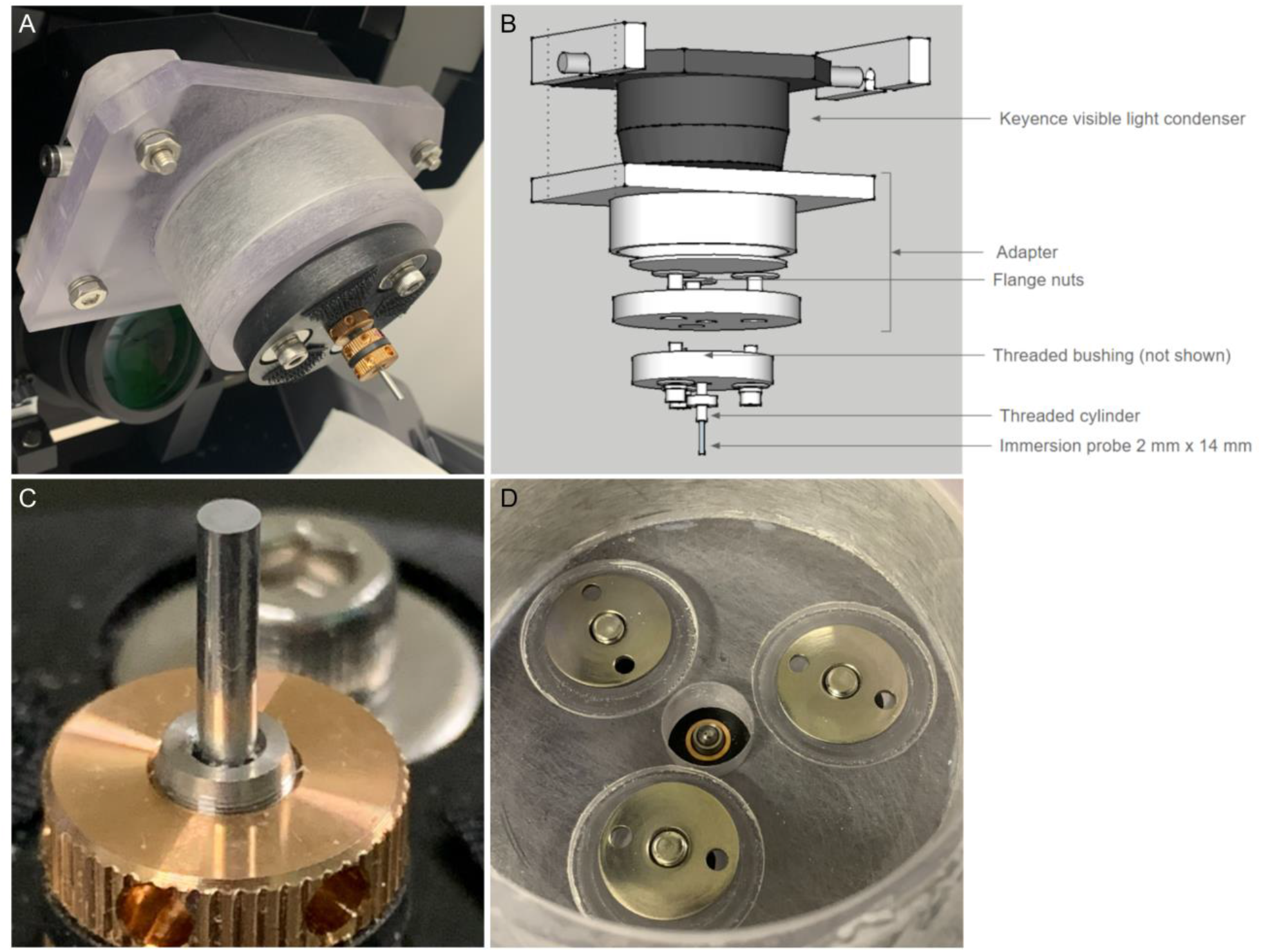
An “immersion probe” regulates buffer depth abovethe sample. An adapter assembled from polycarbonate sheet and cylinder stock is sized to fit over the Keyence visible light condenser unit, clamped to the condenser lateral lift pins by 4 stainless steel cap head bolts. The hinged condenser, tilted up here, is lowered during imaging. (*B*) Exploded diagram of adapter components. A secondary platform, adjustable in the X-Y axis and bolted to the lower surface of the adapter using recessed T-nuts (*D*), holds a finely threaded cylinder (120 turns per inch; 212 µm per turn) turning in a bronze bushing (Thorlabs parts # F19MS100 and F19MSN1P) centered on the optical axis. (*C*) The threaded cylinder supports the immersion probe, a 316 stainless steel cylinder, 2 mm x 14 mm long (McMaster-Carr part #93600A223) polished on the lower face using diamond and aluminum oxide lapping discs (3M-EMSD Diamond Film Type H 5” 3 µm, 3M-EMSD A/O Polish Disc 4” 0.3 µm). The probe is glued into the hexagonal recess of the threaded cylinder using nail polish, which can be dissolved by acetone when there is need to remove it. A pair of knurled adjustment rings (Thorlabs #LN19120) compressing a stiff rubber washer allows manual rotation of the threaded cylinder. A third ring at the base of the cylinder (i.e., adjacent to the secondary platform) serves as a lock nut to stabilize the cylinder in the desired position. When imaging, the lower face of the immersion probe is centered in the microscope optical axis, covers the entire field of view, and is parallel to the plane of focus, typically 30-40 microns above the surface of the sample. The height of the probe remains fixed during the experiment. (*D*) View of bronze bushing and recessed T-nuts in oversized holes in the lower platform allowing adjustment in the X-Y axes.

**FIGURE S2.**
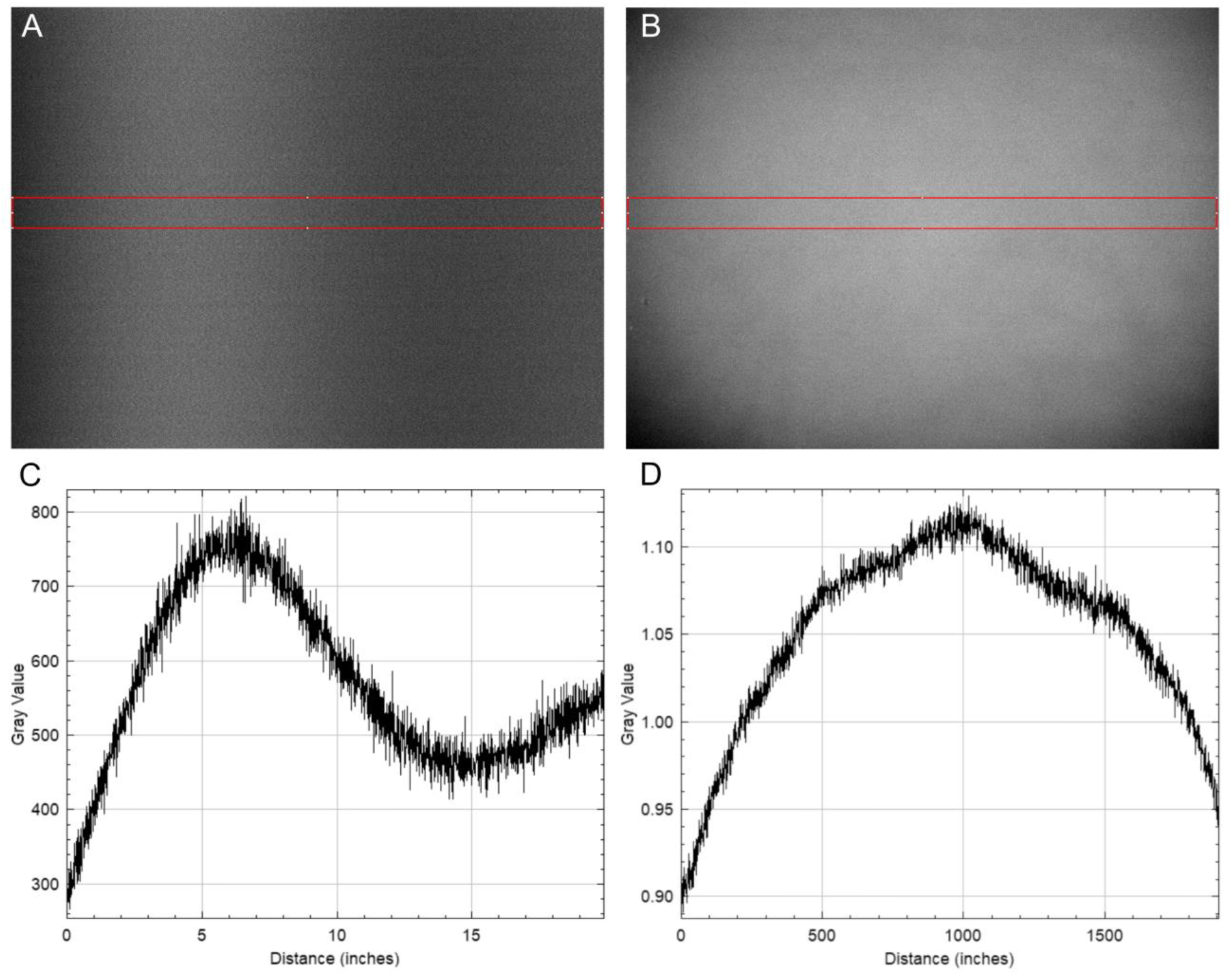
Background correction images. (*A*) “System ghost” and (*B*) relative intensity (“vignetting”) images and their quantification (*C,D*) illustrate variation in signal arising from non-antigen sources in the optical system. Appropriate corrections are made to raw image data to minimize signal from these sources.

**FIGURE S3.**
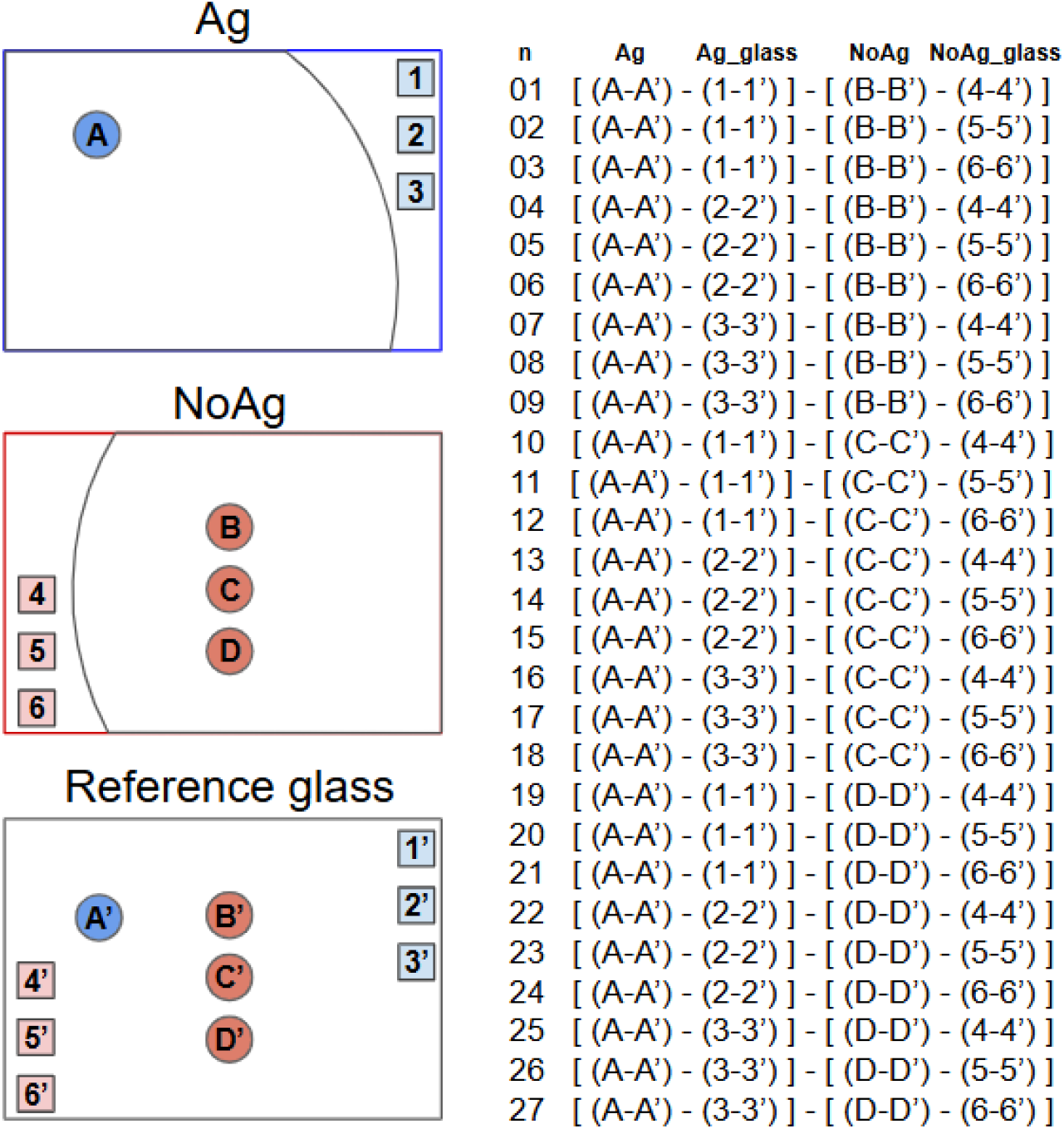
Schematic of image analysis strategy. At each experimental timepoint, six sets of ROIs, two each from three images, contribute to the calculation of Ag-specific signal. An Ag-specific ROI (e.g., circle “A”) and ≥ 3 “*cis*-glass” ROIs (“1-3”) in adjacent buffer regions are defined in the image of the Ag-containing sample (top panel); ≥ 3 NoAg ROIs (“B-D”) and ≥ 3 “*cis*-glass” NoAg ROIs (“4-6”) in adjacent buffer regions are defined in the image of the NoAg-containing negative control sample (middle panel); ROIs from the Ag and NoAg images (“A’-D’” and “1’-6’” ) are quantified in a reference glass image from a region of the coverslip on the same TMA (lower panel). Calculation of Ag-specific raw signal is as shown in the table to the right. NoAg ROIs B-D correct for Ag-independent sample autofluorescence and non-specific Ab binding to the sample. “*Cis*-glass” ROIs in each image correct for local variation in the depth of staining buffer. Reference glass ROIs correct for variation in sample-independent signal in the optical path, primarily contributed by the immersion probe surface. The raw signal for each time point is converted to molar [AgAb] and plotted as the mean ± 2 SD of the replicate calculations.

**FIGURE S4.**
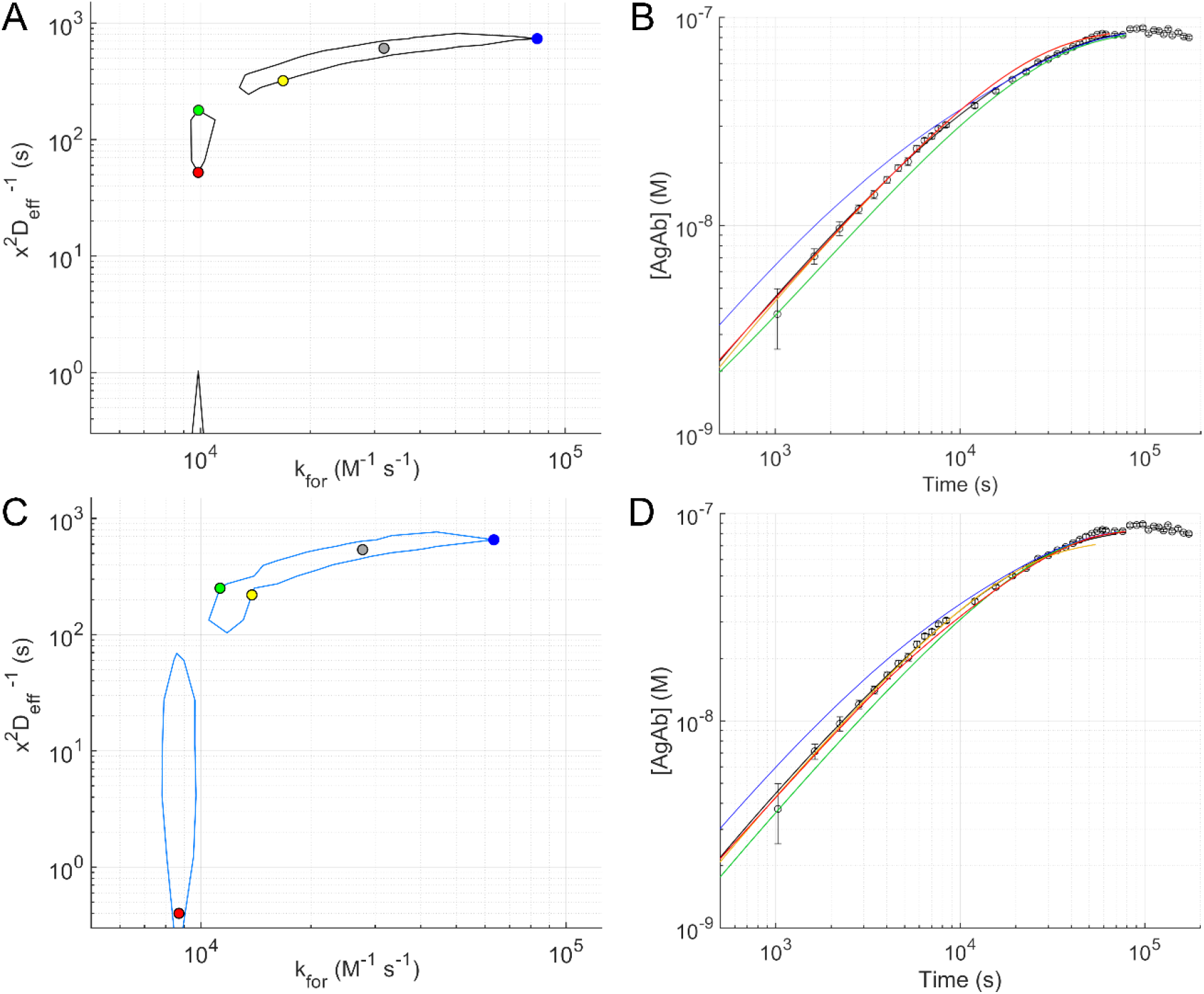
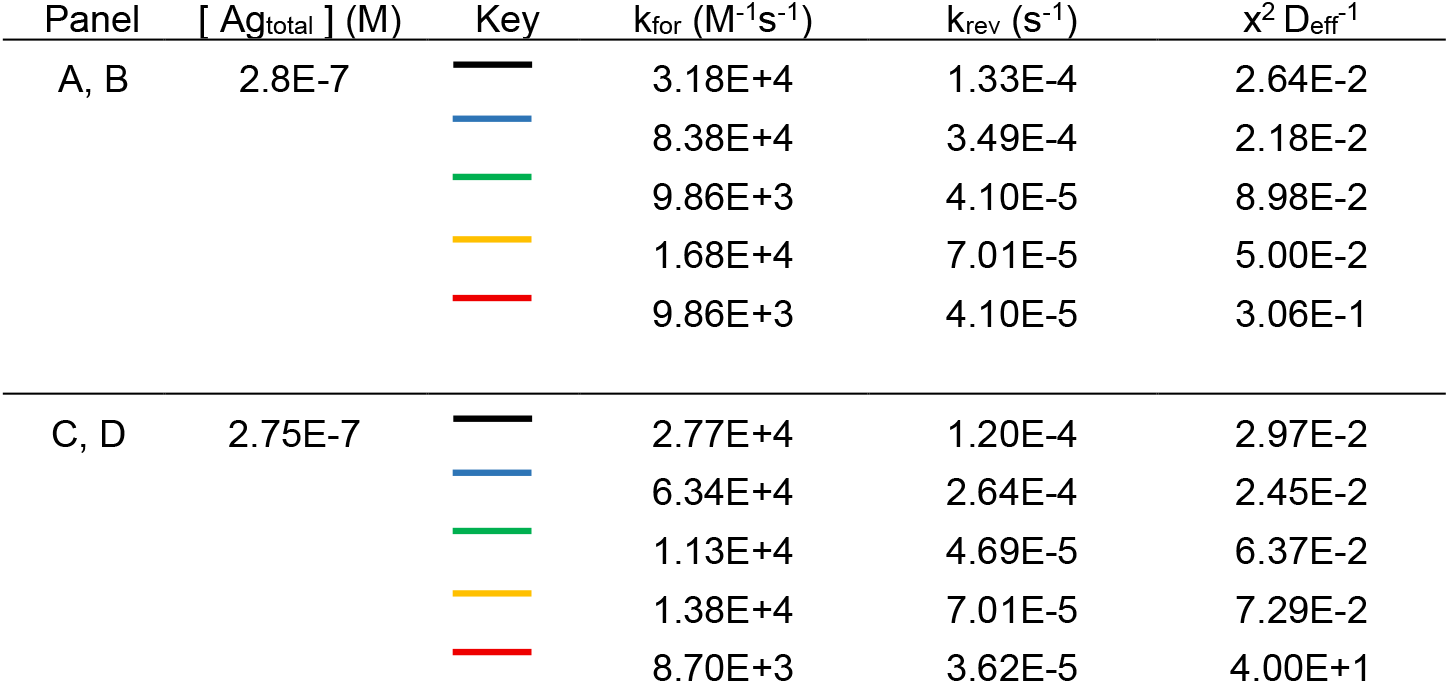
Points on contour plot boundaries correlate with varied reaction time courses. Contour plot perimeters are shown from Fig. 4K ROI 01 (*A*, Nominal fit; C, K_D error_min_). Selected points on the contour plot boundaries are correlated with their corresponding predicted time courses (*B, D*, respectively; Fast_Rx_Dif.exe), which have different slopes and intercepts, bounding the observed experimental data. Corresponding reaction parameters are listed in Figure Table S4, below. **FIGURE TABLE S4**

**FIGURE S5.**
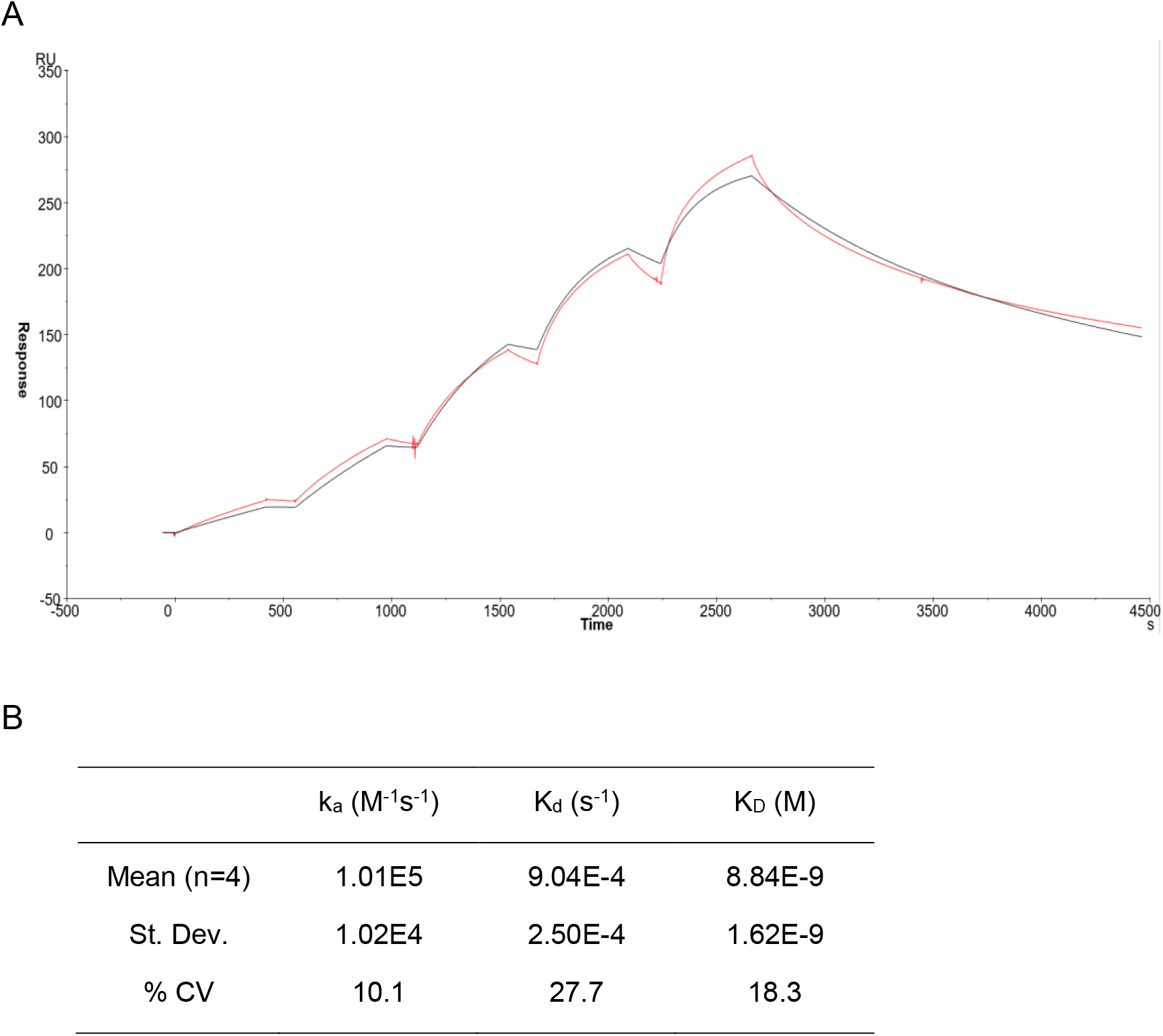
Surface plasmon resonance analysis. (*A*) Sensorgrams for Clone 124 mAb binding to human BCL2 aa 41-54 via single-cycle analysis on Biacore. Clone 124 anti-BCL2 mAb was injected over captured biotinylated BCL2 peptide on a streptavidin chip (Sensor Chip SA) on a Biacore T200 system. Red line corresponds to raw data; black line corresponds to a fit to the bivalent analyte model. Antibody concentrations injected were (in order) 0.74, 2.22, 6.66, 20, and 60 nM. (*B*) Kinetic parameters determined from 4 replicate experiments.

**FIGURE S6.**
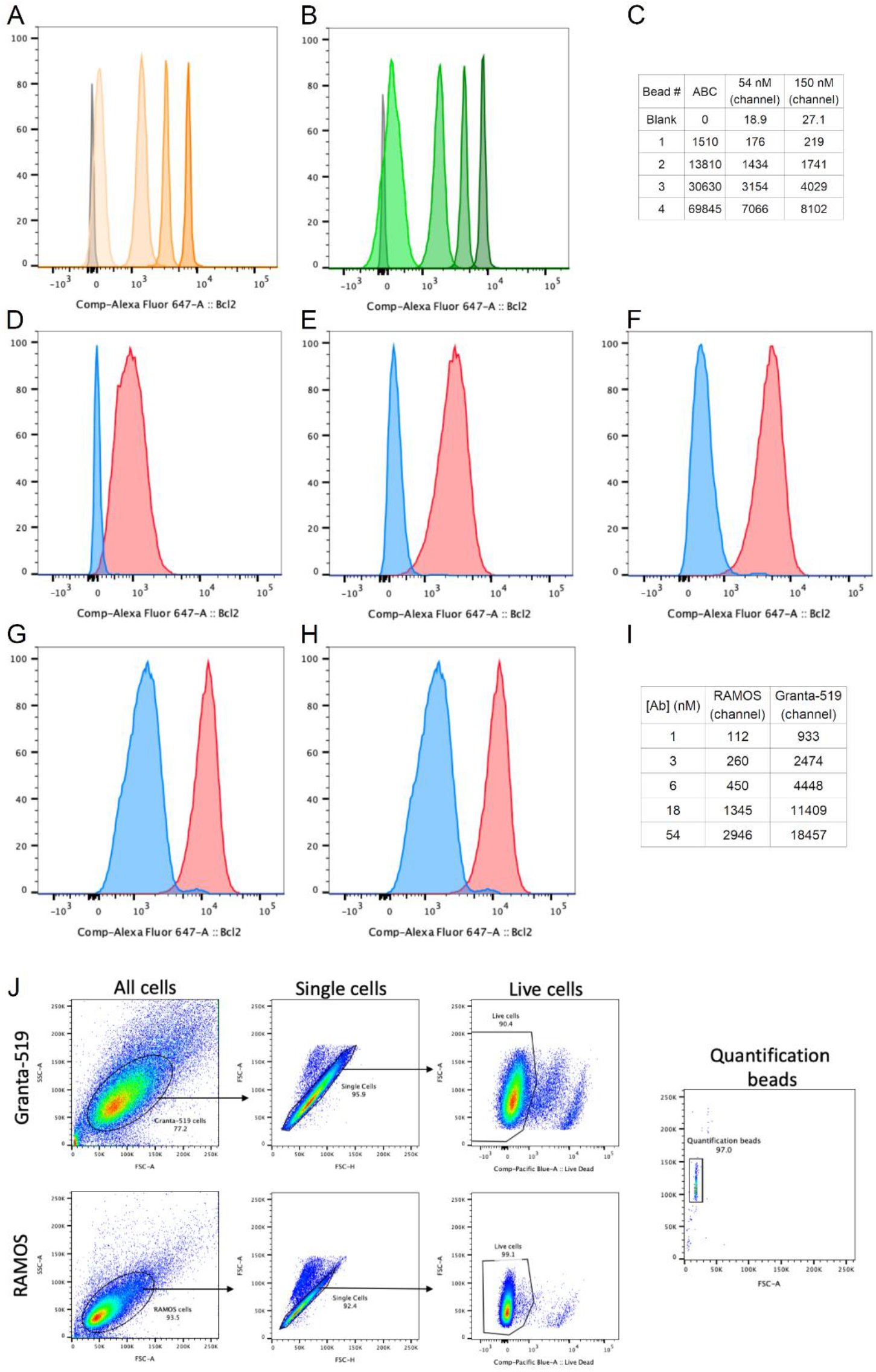
FACS analysis of BCL2 expression in Granta-519 and RAMOS lymphoid cells. (*A, B*) Histograms of Bangs lab QSC calibration beads stained with 54 nM (orange) and 150 nM (green) mouse anti-BCL2 clone 124, respectively, or naive IgG control antibody (grey). (*C*) MFI (channel) values for calibration beads with defined ABC. (*D-H*) Granta-519 (red) and RAMOS (blue) cell stained with anti-BCL2 at (*D*) 1 nM, (*E*) 3 nM, (*F*) 6 nM, (*G*) 18 nM, and (*H*) 54 nM. (*I*) Mean channel MFI (channel) values for Granta and RAMOS cell histograms; cell diameters measured for fixed and permeabilized Granta-519 and RAMOS cells prepared in parallel were 14.0 and 11.0 microns, respectively. (*J*) Gating strategy, representative images.

**FIGURE S7.**
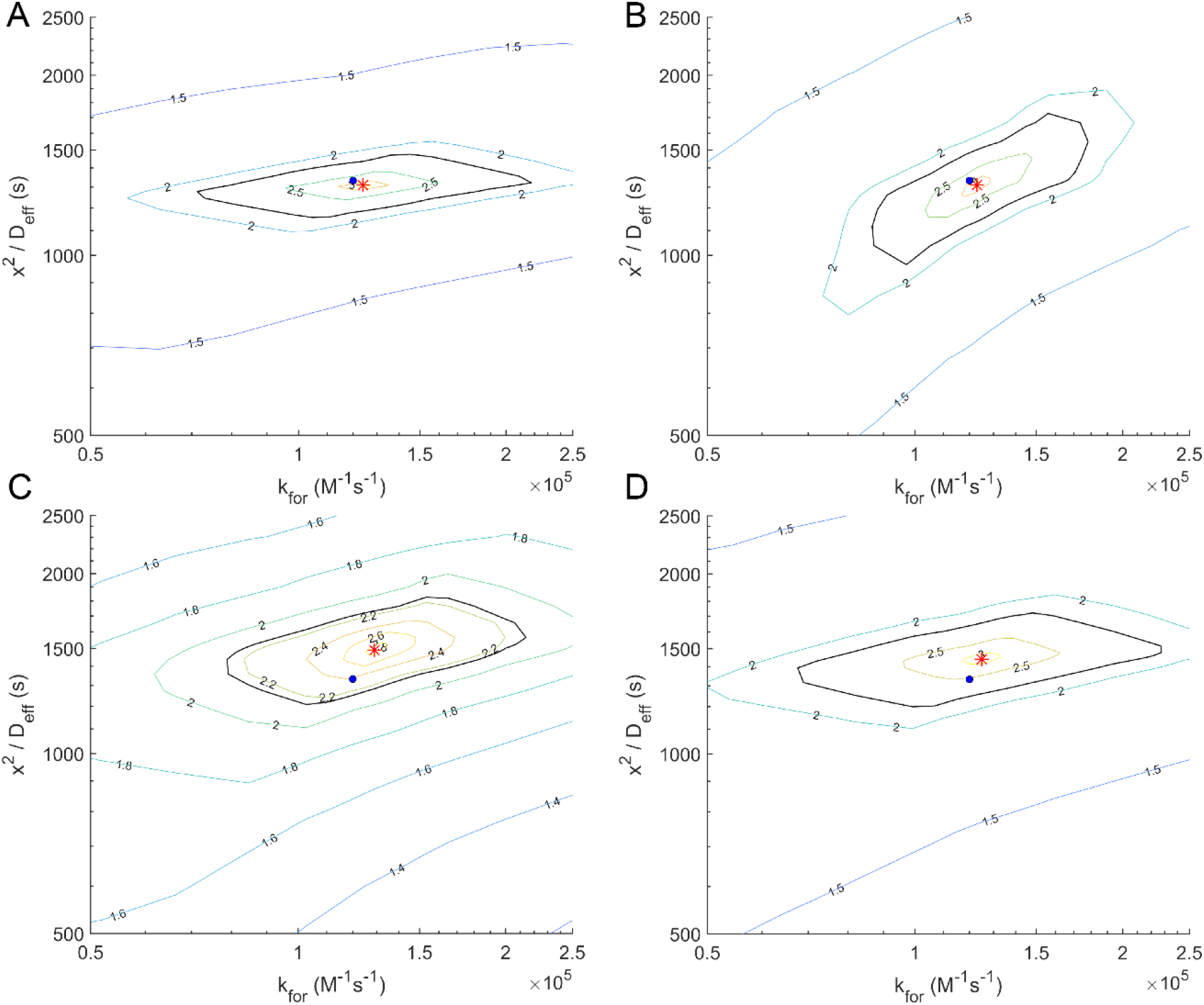
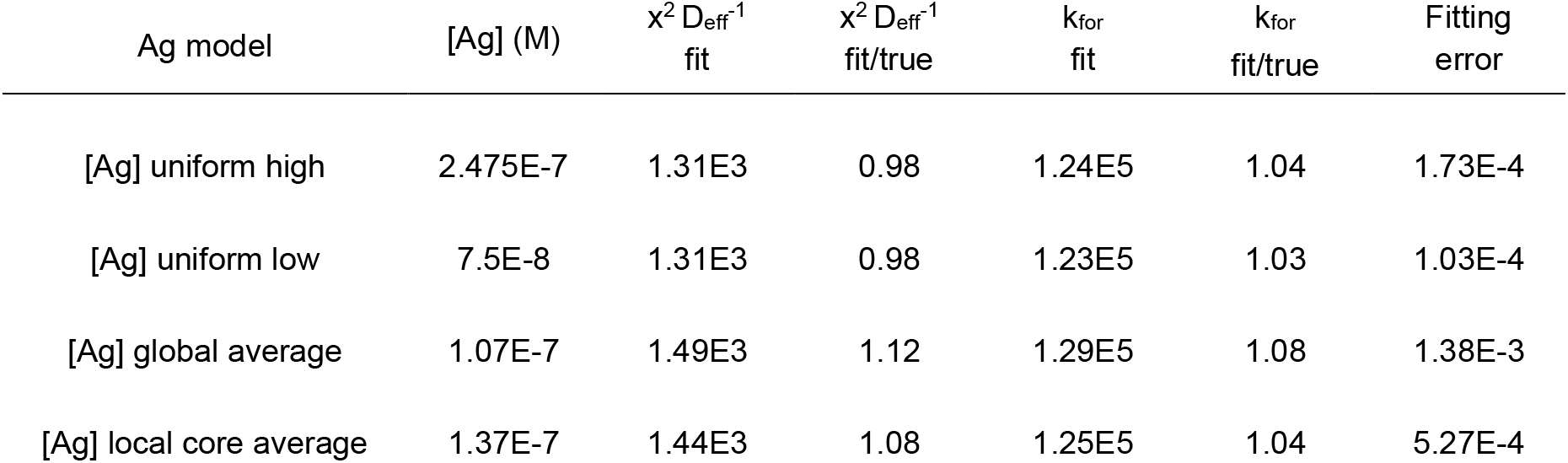
Parameter fitting is within 15% of “true” values for spatially heterogeneous Ag distribution. MATLAB parameter fitting results for Wasserstein mixed-[Ag] models simulated with Virtual Cell. Red asterisk: optimum MATLAB fit parameters; blue marker: expected (*i*.*e*., Virtual Cell model) values for k_for_ and x^2^/D_eff_ (1.2E5 M^-1^sec^-1^ and 1.33E3 sec, respectively). The black contour plot error boundary at 0.7% includes within it the expected values of k_for_ and x^2^/D_eff_ for models with spatially uniform high-[Ag] (*A*), uniform low-[Ag] (*B*), average [Ag] calculated for the entire 25 µm-wide model (*C*) and [Ag] averaged over the central 5 µm-wide high-Ag and flanking 4 µm-wide low-Ag regions (*D*). (see Fig. 8 D-F.) Fitting parameters are summarized in Figure Table S7, below. **FIGURE TABLE S7**

**FIGURE S8.**
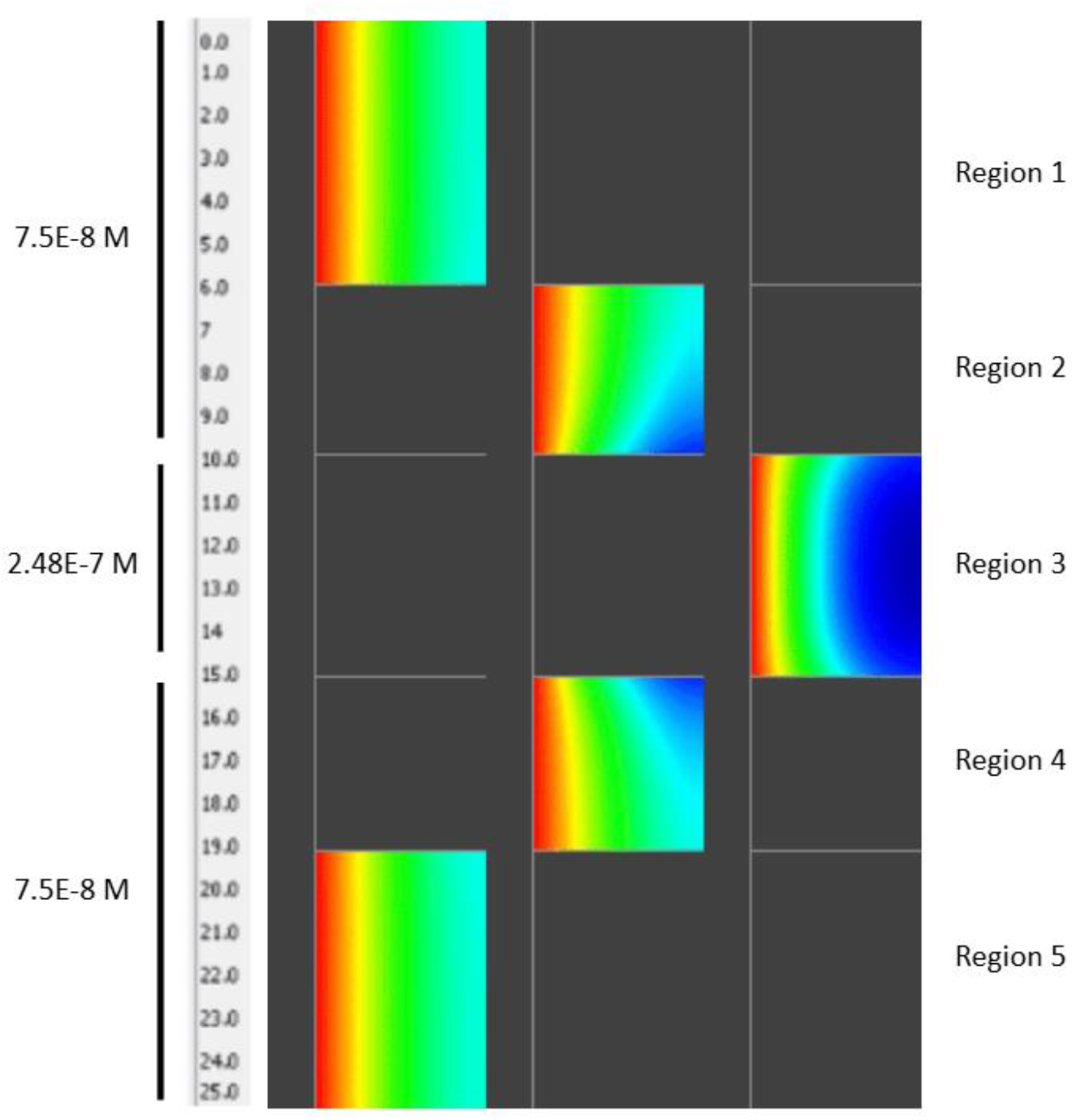
Antigen spatial heterogeneity affects IHC reaction rate. The image above is a still frame from a VCell simulation of AgAb complex formation in a sample with regions of different antigen concentration. Region 3, with 3.3-fold higher [Ag] than the surrounding regions, requires more time to reach equilibrium. Antibody diffusing into region 3 from the adjacent regions 2 and 4 accelerates the formation of AgAb in the margins of region 3, while slowing the formation of AgAb in regions 2 and 4 relative to the rate of formation in the most lateral regions 1 and 5. See separate Fig_S8.gif file for full simulation. Despite these interactions, the IHC reaction parameters in the sample can be determined to within 15% of their true values.

**FIGURE S9.**
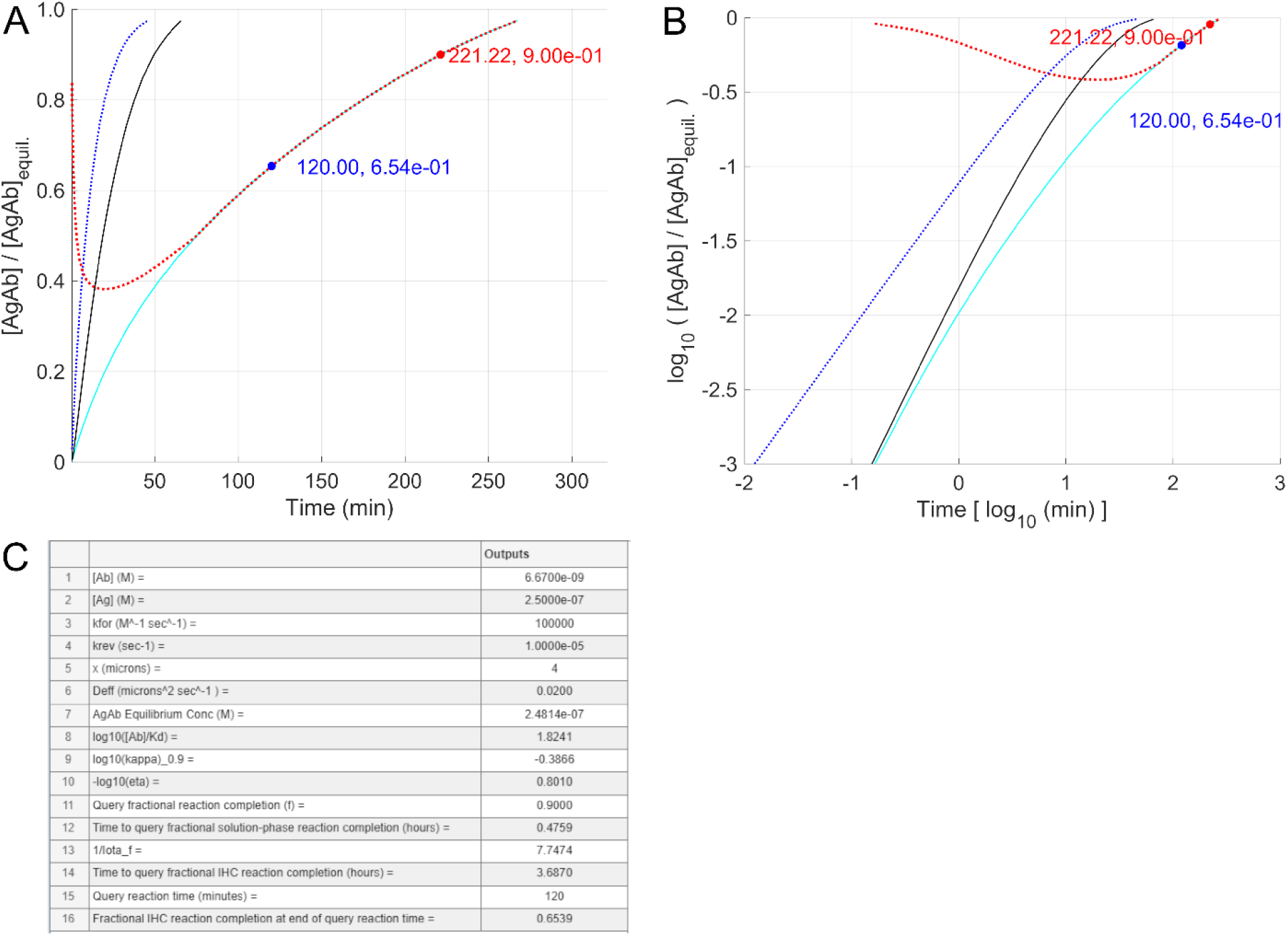
Dynamic range compression. “Dynamic range compression” is the extent to which a comparison of the [AgAb] yield for two reactions at the same timepoint underestimates the relative difference in [AgAb] that would be observed when both reactions reach equilibirium. The upper two figures show the time courses on linear (*A*) and log (*B*) scales for a primary [Ag] (2.5E-7 M; cyan) and 10-fold lower reference [Ag] (2.5E-8 M; black). Other reaction parameters: [Ab] = 6.67E-9 M (1 µg/mL); k_for_ = 1E5 M^-1^s^-1^; k_rev_ = 1E-5 s^-^1; tissue depth = 4 µm; D_eff_ = 2E-2 µm^2^s^-1^. The dotted red curve shows the dynamic range compression (cyan vs. black) at different times. Blue dotted curves represent the time course for solution-phase reactions with the same [Ab], k_for_ and k_rev_ parameters. Blue text indicates 60 minutes reaction time and the corresponding fractional reaction progress towards equilibrium. The red text indicates the time (min) needed to reach 90% of reaction completion. (*C*) The table summarizes the relevant reaction parameters and model predictions. Data are from Fast_Rx_dif_model.exe.

## SUPPLEMENTAL TABLES

**TABLE S1.** Recommended IHC antibody on-slide concentrations vary > 150-fold. The recommended on-slide antibody concentrations for 30 commercial products range from 5E-10 M to 7.8E-8 M. Molar concentrations assume antibody mass is 1.5E5 Da. URLs for data sources:

| Antibody | Stock concentration<br>($\mu\text{g/mL}$) | On-slide concentration<br>($\mu\text{g/mL}$) | On-slide concentration<br>(M) |
| --- | --- | --- | --- |
| 1. BOND anti-Desmoglein 3 (EPR14101) | 11.7 | 11.7 | 7.80E-8 |
| 2. BOND anti-CD4 (4B12) | 6.6 | 6.6 | 4.40E-8 |
| 3. CONFIRM anti-BCL2 (124) | 17 | 4.25 | 2.80E-8 |
| 4. CONFIRM anti-CD15 (MMA) | 11 | 2.8 | 1.90E-8 |
| 5. BOND anti-ALK (5A4) | 2.6 | 2.6 | 1.70E-8 |
| 6. BOND anti-SOX10 (SOX10/991) | 2 | 2 | 1.30E-8 |
| 7. BOND anti-IgG4 (HP6023) | 2 | 2 | 1.30E-8 |
| 8. BOND anti-Ki67 (MM1) | 1.9 | 1.9 | 1.30E-8 |
| 9. CONFIRM anti-PD-L1 (SP142) | 7.2 | 1.8 | 1.20E-8 |
| 10. PATHWAY anti-Her2 (4B5) | 6 | 1.5 | 1.00E-8 |
| 11. CONFIRM anti-CD10 (SP67) | 4.9 | 1.23 | 8.20E-9 |
| 12. BOND anti-PR (16) | 1 | 1 | 6.70E-9 |
| 13. BOND anti-EpCAM (EPR20532-222) | 1 | 1 | 6.70E-9 |
| 14. BOND anti-GLUT1 (GLUT1/2475) | 1 | 1 | 6.70E-9 |
| 15. BOND anti-CD3 (LN10) | 1 | 1 | 6.70E-9 |
| 16. BOND anti-LEF-1 (EPR2029Y) | 0.9 | 0.9 | 6.00E-9 |
| 17. BOND anti-ER (6F11) | 0.9 | 0.9 | 5.90E-9 |
| 18. CONFIRM anti-CD4 (SP35) | 2.5 | 0.625 | 4.20E-9 |
| 19. BOND anti-Bcl-2 (bcl-2/100/D5) | 0.5 | 0.5 | 3.30E-9 |
| 20. CONFIRM anti-PD-L1 (SP263) | 1.6 | 0.4 | 2.70E-9 |
| 21. BOND anti-H-caldesmon (BSB-19) | 0.4 | 0.4 | 2.70E-9 |
| 22. BOND anti-Arginase-1 (EPR6672(B)) | 0.3 | 0.3 | 2.00E-9 |
| 23. BOND anti-CD8 (4B11) | 0.25 | 0.25 | 1.70E-9 |
| 24. CONFIRM anti-ER (SP1) | 1 | 0.25 | 1.70E-9 |
| 25. CONFIRM anti-PR (1E2) | 1 | 0.25 | 1.70E-9 |
| 26. BOND anti-c-erbB-2 (CB11) | 0.15 | 0.15 | 1.00E-9 |
| 27. CONFIRM anti-CD68 (KP-1) | 0.4 | 0.1 | 6.70E-10 |
| 28. CONFIRM anti-CD8 (SP57) | 0.4 | 0.1 | 6.70E-10 |
| 29. CONFIRM anti-CD20 (L26) | 0.3 | 0.075 | 5.00E-10 |
| 30. CONFIRM anti-CD79a (SP18) | 0.3 | 0.075 | 5.00E-10 |

1. https://files.leicabiosystems.com/LBS/US/All/en?keycode=PA0347-U
2. https://files.leicabiosystems.com/LBS/US/All?keycode=PA0427&lot-range=PA0427+BP
3. https://elabdoc-prod.roche.com/eLD/api/downloads/53aebeee-0c15-ee11-2091-005056a71a5d?countryIsoCode=us
4. https://elabdoc-prod.roche.com/eLD/api/downloads/9baf2727-7233-ea11-fa90-005056a772fd?countryIsoCode=us
5. https://files.leicabiosystems.com/LBS/US/All/en?keycode=PA0831&lot-range=PA0831-BP
6. https://files.leicabiosystems.com/LBS/US/All/en?keycode=PA0010-U
7. https://files.leicabiosystems.com/LBS/US/All/en?keycode=PA0977-U
8. https://files.leicabiosystems.com/LBS/US/All?keycode=PA0118&lot-range=PA0118
9. https://elabdoc-prod.roche.com/eLD/api/downloads/5c519e5b-b653-ec11-0f91-005056a71a5d?countryIsoCode=XG
10. https://elabdoc-prod.roche.com/eLD/api/downloads/fc569ff5-2236-ea11-fc90-005056a71a5d?countryIsoCode=us
11. https://elabdoc-prod.roche.com/eLD/api/downloads/eba02455-b334-ea11-fa90-005056a772fd?countryIsoCode=u
12. https://files.leicabiosystems.com/LBS/US/All?keycode=PA0436&lot-range=PA0436
13. https://files.leicabiosystems.com/LBS/US/All?keycode=PA0310-U
14. https://files.leicabiosystems.com/LBS/US/All?keycode=PA0866-U
15. https://files.leicabiosystems.com/LBS/US/All?keycode=PA0122&lot-range=PA0122
16. https://files.leicabiosystems.com/LBS/US/All?keycode=PA0756-U
17. https://files.leicabiosystems.com/LBS/US/All?keycode=PA0151-U&lot-range=PA0151-U
18. https://elabdoc-prod.roche.com/eLD/api/downloads/744ae8e1-1613-ea11-fa90-005056a772fd?countryIsoCode=us
19. https://files.leicabiosystems.com/LBS/US/All?keycode=PA0117&lot-range=PA0117
20. https://elabdoc-prod.roche.com/eLD/api/downloads/eba02455-b334-ea11-fa90-005056a772fd?countryIsoCode=us
21. https://files.leicabiosystems.com/LBS/US/All?keycode=PA0713-U
22. https://files.leicabiosystems.com/LBS/US/All?keycode=PA0101-U
23. https://files.leicabiosystems.com/LBS/US/All?keycode=PA0183&lot-range=PA0183+BP
24. https://elabdoc-prod.roche.com/eLD/api/downloads/db5da765-49fa-ed11-1d91-005056a71a5d?countryIsoCode=us
25. https://elabdoc-prod.roche.com/eLD/api/downloads/76ea4fea-e112-ea11-fa90-005056a772fd?countryIsoCode=us
26. https://files.leicabiosystems.com/LBS/US/All?keycode=PA0983-U&lot-range=PA0983-U+Rev+B
27. https://elabdoc-prod.roche.com/eLD/api/downloads/e8e0fa52-6e33-ea11-fa90-005056a772fd?countryIsoCode=us
28. https://elabdoc-prod.roche.com/eLD/api/downloads/55085a38-67c7-ed11-1a91-005056a71a5d?countryIsoCode=us
29. https://elabdoc-prod.roche.com/eLD/api/downloads/50d4b69b-6833-ea11-fa90-005056a772fd?countryIsoCode=us
30. https://elabdoc-prod.roche.com/eLD/api/downloads/f693e26c-8d34-ea11-fa90-005056a772fd?countryIsoCode=us

**TABLE S2.** Cr_*f*_ values. *Cr*_*f*_ relates null tissue diffusion time (from Eq. 1) to x^2^/D_eff_ as noted in Eq. 9.

| $f$ | $Cr_f$ | $f$ | $Cr_f$ | $f$ | $Cr_f$ | $f$ | $Cr_f$ | $f$ | $Cr_f$ | $f$ | $Cr_f$ |
| --- | --- | --- | --- | --- | --- | --- | --- | --- | --- | --- | --- |
| 0.99 | 5.614E-1 | 0.79 | 1.827E+0 | 0.59 | 3.618E+0 | 0.39 | 8.371E+0 | 0.19 | 3.527E+1 | 0.0075 | 2.268E+4 |
| 0.98 | 6.665E-1 | 0.78 | 1.892E+0 | 0.58 | 3.750E+0 | 0.38 | 8.818E+0 | 0.18 | 3.930E+1 | 0.005 | 5.115E+4 |
| 0.97 | 7.484E-1 | 0.77 | 1.959E+0 | 0.57 | 3.888E+0 | 0.37 | 9.301E+0 | 0.17 | 4.406E+1 | 0.0035 | 1.048E+5 |
| 0.96 | 8.200E-1 | 0.76 | 2.027E+0 | 0.56 | 4.033E+0 | 0.36 | 9.824E+0 | 0.16 | 4.974E+1 | 0.0025 | 2.073E+5 |
| 0.95 | 8.857E-1 | 0.75 | 2.098E+0 | 0.55 | 4.186E+0 | 0.35 | 1.039E+1 | 0.15 | 5.659E+1 | 0.0015 | 5.944E+5 |
| 0.94 | 9.478E-1 | 0.74 | 2.170E+0 | 0.54 | 4.346E+0 | 0.34 | 1.101E+1 | 0.14 | 6.496E+1 | 0.001 | 1.432E+6 |
| 0.93 | 1.007E+0 | 0.73 | 2.244E+0 | 0.53 | 4.516E+0 | 0.33 | 1.169E+1 | 0.13 | 7.534E+1 |  |  |
| 0.92 | 1.066E+0 | 0.72 | 2.321E+0 | 0.52 | 4.694E+0 | 0.32 | 1.243E+1 | 0.12 | 8.843E+1 |  |  |
| 0.91 | 1.123E+0 | 0.71 | 2.400E+0 | 0.51 | 4.883E+0 | 0.31 | 1.325E+1 | 0.11 | 1.052E+2 |  |  |
| 0.9 | 1.179E+0 | 0.7 | 2.482E+0 | 0.5 | 5.083E+0 | 0.3 | 1.415E+1 | 0.1 | 1.273E+2 |  |  |
| 0.89 | 1.235E+0 | 0.69 | 2.567E+0 | 0.49 | 5.295E+0 | 0.29 | 1.514E+1 | 0.09 | 1.572E+2 |  |  |
| 0.88 | 1.292E+0 | 0.68 | 2.655E+0 | 0.48 | 5.520E+0 | 0.28 | 1.624E+1 | 0.08 | 1.989E+2 |  |  |
| 0.87 | 1.348E+0 | 0.67 | 2.745E+0 | 0.47 | 5.759E+0 | 0.27 | 1.747E+1 | 0.07 | 2.598E+2 |  |  |
| 0.86 | 1.405E+0 | 0.66 | 2.840E+0 | 0.46 | 6.013E+0 | 0.26 | 1.883E+1 | 0.06 | 3.537E+2 |  |  |
| 0.85 | 1.463E+0 | 0.65 | 2.938E+0 | 0.45 | 6.285E+0 | 0.25 | 2.037E+1 | 0.05 | 5.093E+2 |  |  |
| 0.84 | 1.521E+0 | 0.64 | 3.039E+0 | 0.44 | 6.574E+0 | 0.24 | 2.211E+1 | 0.04 | 7.958E+2 |  |  |
| 0.83 | 1.580E+0 | 0.63 | 3.145E+0 | 0.43 | 6.884E+0 | 0.23 | 2.407E+1 | 0.03 | 1.415E+3 |  |  |
| 0.82 | 1.640E+0 | 0.62 | 3.256E+0 | 0.42 | 7.217E+0 | 0.22 | 2.631E+1 | 0.02 | 3.184E+3 |  |  |
| 0.81 | 1.701E+0 | 0.61 | 3.371E+0 | 0.41 | 7.573E+0 | 0.21 | 2.887E+1 | 0.015 | 5.661E+3 |  |  |
| 0.8 | 1.763E+0 | 0.6 | 3.492E+0 | 0.4 | 7.957E+0 | 0.2 | 3.183E+1 | 0.01 | 1.275E+4 |  |  |

**TABLE S3.** Representative imaging sequence. Representative imaging time points are described for sample incubation with 3 successive antibody concentrations. Imaging intervals balance sampling frequency against photobleaching.

| [Ab] (M) | Time course | Interval (min) | Cycles | Interval Duration (hr) | Cummulative Duration (hr) |
| --- | --- | --- | --- | --- | --- |
| 1E-9 | TC01 | 5 | 15 | 1.2 | 1.2 |
|  | TC02 | 15 | 10 | 2.5 | 3.7 |
|  | TC03 | 60 | 12 | 12 | 15.7 |
|  | TC04 | 180 | 12 | 36 | 51.7 |
| 3E-9 | TC05 | 6 | 15 | 1.5 | 53.2 |
|  | TC06 | 60 | 17 | 17 | 70.2 |
| 1E-8 | TC07 | 6 | 15 | 1.5 | 71.7 |
|  | TC08 | 60 | 17 | 17 | 88.7 |

**TABLE S4.**
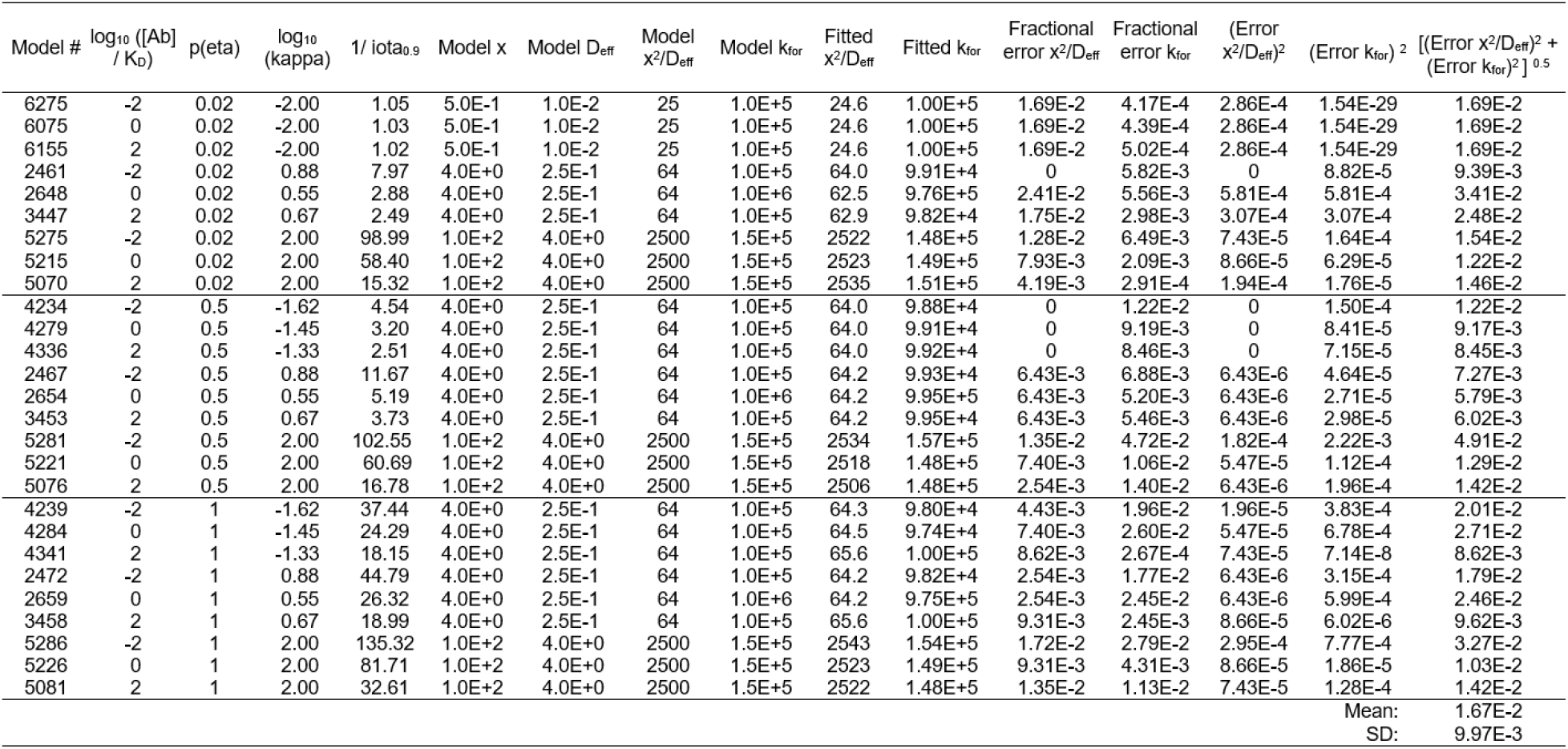
Results of fitting Virtual Cell model reactions with fast interpolation method. Twenty-seven Virtual Cell reaction-diffusion simulations (“Model #”) were run according to the parameters shown in columns 2-9. The resulting time course data were fit by rapid interpolation (see Methods) to yield the optimum x^2^/D_eff_ and k_for_ parameters shown in columns 10, 11. Mean fractional fitting errors (columns 12, 13) were used to calculate individual and global average root mean square errors (column 16).

**TABLE S5.** BSA gel and cell line LOB/LOD data **See separate Excel spreadsheet**.

**TABLE S6.** Estimated reaction parameters for Figure 4 AgAb binding reactions. Fitted reaction parameters for the Ag-BSA gel experimental time courses shown in Figure 4. *A*) ROI 01 and *B*) ROI 12. Left to right: AgAb1-AgAb3, plateau [AgAb] for successively increasing [Ab]; [Ab1] = 9.1E-10 M, [Ab2] = 2.5E-9 M, [Ab3] = 9.8E-9 M; K_D_ mean, [Ag]_total_ and K_D_ error (from MATLAB lest squares fit), calculated from Langmuir equilibrium using [AgAb1] to [AgAb3] and [Ab1] to [Ab3]; x^2^/D_eff_, k_for_, k_rev_ and minimum residual error, best fit values from curve fitting calculations. Rows 1-6, “nominal” to “K_D_ err_min”, are for curve fits resulting from tested plateau [AgAb] values ± 2 SD from the nominal means. Results for K_D max_ and [Ag]_max_ are identical. Mean and SD are based on the 5 unique values in each column. Results for ROI 2-11 are summarized in a separate spreadsheet (Table_S6.xlsx).

| Data | [AgAb1]<br>(M) | [AgAb2]<br>(M) | [AgAb3]<br>(M) | K<sub>D</sub> mean<br>(M) | [Ag]<sub>total</sub><br>(M) | K<sub>D</sub> error | x<sup>2</sup>/D<sub>eff</sub><sup>-1</sup><br>(s) | k<sub>for</sub><br>(M<sup>-1</sup>s<sup>-1</sup>) | k<sub>rev</sub><br>(s<sup>-1</sup>) | minimum<br>error |
| --- | --- | --- | --- | --- | --- | --- | --- | --- | --- | --- |
| nominal | 8.50E-8 | 1.55E-7 | 2.28E-7 | 4.13E-9 | 2.80E-7 | 7.48E-4 | 6.03E+2 | 3.20E+4 | 1.32E-4 | 7.86E-3 |
| K<sub>D</sub> min | 9.11E-8 | 1.53E-7 | 2.23E-7 | 3.41E-9 | 2.61E-7 | 2.48E-4 | 7.35E+2 | 4.24E+4 | 1.45E-4 | 9.73E-3 |
| K<sub>D</sub> max | 7.89E-8 | 1.64E-7 | 2.33E-7 | 5.53E-9 | 3.21E-7 | 3.15E-3 | 4.12E+2 | 1.88E+4 | 1.04E-4 | 8.25E-3 |
| Ag min | 9.11E-8 | 1.47E-7 | 2.23E-7 | 3.43E-9 | 2.58E-7 | 1.26E-3 | 7.42E+2 | 4.14E+4 | 1.42E-4 | 9.74E-3 |
| Ag max | 7.89E-8 | 1.64E-7 | 2.33E-7 | 5.53E-9 | 3.21E-7 | 3.15E-3 | 4.12E+2 | 1.88E+4 | 1.04E-4 | 8.25E-3 |
| K<sub>D</sub> err_min | 8.96E-8 | 1.54E-7 | 2.25E-7 | 3.61E-9 | 2.67E-7 | 2.73E-7 | 7.00E+2 | 3.91E+4 | 1.41E-4 | 6.94E-3 |
| Mean | 8.72E-8 | 1.54E-7 | 2.26E-7 | 4.20E-9 | 2.77E-7 | 1.08E-3 | 6.39E+2 | 3.48E+4 | 1.33E-4 | 8.50E-3 |
| SD. | 5.23E-9 | 6.09E-9 | 4.29E-9 | 8.90E-10 | 2.61E-8 | 1.26E-3 | 1.38E+2 | 9.81E+3 | 1.68E-5 | 1.22E-3 |

| Data | [AgAb1]<br>(M) | [AgAb2]<br>(M) | [AgAb3]<br>(M) | K<sub>D</sub> mean<br>(M) | [Ag]<sub>total</sub><br>(M) | K<sub>D</sub> error | x<sup>2</sup>/D<sub>eff</sub><sup>-1</sup><br>(s) | k<sub>for</sub><br>(M<sup>-1</sup>s<sup>-1</sup>) | k<sub>rev</sub><br>(s<sup>-1</sup>) | minimum<br>error |
| --- | --- | --- | --- | --- | --- | --- | --- | --- | --- | --- |
| nominal | 5.12E-8 | 9.32E-8 | 1.42E-7 | 4.35E-9 | 1.74E-7 | 5.39E-4 | 1.49E+3 | 2.11E+4 | 9.16E-5 | 1.53E-2 |
| K<sub>D</sub> min | 5.50E-8 | 9.32E-8 | 1.40E-7 | 3.71E-9 | 1.66E-7 | 8.05E-4 | 1.71E+3 | 2.53E+4 | 9.38E-5 | 1.15E-2 |
| K<sub>D</sub> max | 4.75E-8 | 9.93E-8 | 1.43E-7 | 5.75E-9 | 1.99E-7 | 5.23E-3 | 1.27E+3 | 1.60E+4 | 9.20E-5 | 1.05E-2 |
| Ag min | 5.50E-8 | 8.71E-8 | 1.40E-7 | 3.78E-9 | 1.64E-7 | 3.38E-3 | 1.75E+3 | 2.51E+4 | 9.48E-5 | 1.15E-2 |
| Ag max | 4.75E-8 | 9.93E-8 | 1.43E-7 | 5.75E-9 | 1.99E-7 | 5.23E-3 | 1.27E+3 | 1.60E+4 | 9.20E-5 | 1.05E-2 |
| K<sub>D</sub> err_min | 5.26E-8 | 9.32E-8 | 1.41E-7 | 4.08E-9 | 1.71E-7 | 2.59E-7 | 1.53E+3 | 2.18E+4 | 8.91E-5 | 1.37E-2 |
| Mean | 5.23E-8 | 9.32E-8 | 1.41E-7 | 4.33E-9 | 1.75E-7 | 1.99E-3 | 1.55E+3 | 2.19E+4 | 9.23E-5 | 1.25E-2 |
| SD | 3.09E-9 | 4.31E-9 | 1.04E-9 | 8.30E-10 | 1.43E-8 | 2.23E-3 | 1.91E+2 | 3.78E+3 | 2.20E-6 | 1.94E-3 |

**TABLE S7.** Granta-519 cell parameter fitting data. See separate Excel spreadsheet for ROIs 101-108.

## SUPPLEMENTAL METHODS

### Supporting Methods

#### Parameter estimation from experimental data

Data from 2294 MATLAB-simulated reaction time courses were organized to allow rapid interpolation for other reaction conditions. Reaction data are summarized in a matrix with 6 columns: 1) log_10_([Ab]/K_D_); 2) “kappa_root” = (x^2^/D_eff_)*(2*[Ab]*k_for_ + k_rev_); 3) p(*eta*)= -log_10_(*eta*); 4) log_10_(k_eff_) = log_10_(2*[Ab]*k_for_ + k_rev_); 5) log_10_(seconds); 6) log_10_(fractional reaction completion, *f*). Each time course is described in ∼100 to 250 rows, according to the reaction duration, from *f <* 0.001 to > 0.9. A second matrix contains data for antibody diffusion into a planar volume in the absence of binding, *f* vs. *Cr* (Table S2). Using these data, the program calculates, for 28 representative values of *f =* 0.001 to 0.95, the value of *iota*_*f*_ (Eq. 10) and *kappa*_*f*_ (Eq. 11,12) for each of the modeled reactions.

In a 3-dimensional space defined by the unitless parameters log_10_([Ab]/K_D_), log_10_(*kappa*_*f*_) and p(*eta*) = -log_10_(*eta*), the modeled reactions lie in a regular grid, where each point in the grid is associated with a single value of *iota*_*f*_. The MATLAB “scatteredInterpolant” function uses the known value of *iota*_*f*_ at each point in the parameter space to interpolate the value of *iota*_*f*_ for the user-input reaction parameters. The interpolated values of *iota*_*f*_ are then used, with Eq. 10, to calculate the IHC reaction time (IHC reaction time = solution-phase reaction time / iota_*f*_). The model data set support interpolation from *f < 0*.*001* through *f* = 0.9; results for *f* > 0.9 will be variably less reliable.

The curve fitting calculations require values for [Ab]_1-3_, section thickness (*x*), [Ag]_total_ and K_D_, along with experimental time course data expressed as *f* vs. time (s). The output of the curve fitting are a set of x^2^/D_eff_ and k_for_ parameter pairs consistent with the experimental time course data.

Antibody concentrations [Ab]_1-3_ are controlled by the investigator, but to account for experimental variation from the intended values, the observed specific fluorescence was measured at all time points in reference glass images, corrected by subtracting signal from antibody diluent buffer. The resulting data (intended [Ab] vs. observed specific fluorescence at each [Ab]) was used in least squares linear fitting to estimate the actual [Ab] in each Ab solution, and those values were used to establish the antibody specific fluorescence (fluorescence intensity / sec*M*µm buffer depth).

The total antigen concentration and antibody-antigen K_D_ were estimated using MATLAB least squares fitting according to the Langmuir equilibrium (Eq. 2), using as inputs [Ab]_1_-_3_ (calculated as described above) and [AgAb]_equil_1_ - _3_. To estimate [AgAb]_equil_1-3_, a moving mean with a symmetrical window spanning 5 data points was scanned across the non-zero [AgAb] data at each [Ab]. The time point of maximum value was noted as t_equil. [AgAb]_equil_ mean and standard deviation were calculate using the non-zero values from timepoints at t_equil - 2 (to account for the window size of 5) to the last time point in the [Ab]. In the case where t_equil was less than 3 time points from the final time point, the last 5 non-zero time points were used for the calculations.

Experimental variability in the measured [AgAb]_equil-1-3_ allows a range of possible [Ag]_total_ and K_D_ values. To account for this, the Langmuir calculation was done iteratively, combinatorially scanning [AgAb]_equil-1-3_ values across a range ± 2 standard deviations from the means. The resulting [Ag]_total_ and K_D_ values were sorted to identify 6 sets of reaction parameters, plotted in contour plots with corresponding borders color / patterns as follows: 1) “nominal” (black solid), based on the observed equilibrium [AgAb] at each [Ab]; 2) K_D_min_ (black dash-dot) and 3) K_D_max_ (black dashed), each based on the [AgAb] values yielding the minimum and maximum K_D_; 4) [Ag]_min_ (black dotted) and 5) [Ag]_max_ (black dashed), each based on the [AgAb] values yielding the minimum and maximum [Ag]_total_; 6) fitting error_min_ (solid blue), based on the [AgAb] values yielding the K_D_ with the minimum least squares fitting error for the Langmuir equilibrium. In practice, the [Ag]_max_ and K_D_max_ parameters were typically identical. For each ROI, each of these 5 or 6 sets of unique [Ag]_total_ and K_D_ parameters were used in the curve fitting steps below.

Using pairs of [Ag]_total_ and K_D_ values calculated as described above, 3190 combinations of x^2^/D_eff_ and k_for_ are tested to find those most consistent with the observed time to 50% reaction completion (or another percent reaction completion value, as specified by the “midpoint” parameter). To do this, experimental IHC time point data, x=time (s), are normalized to parameter-less values:

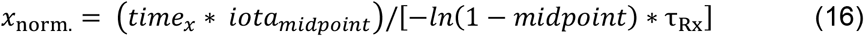

The denominator in this equation is the solution-phase time to the fractional reaction completion specified by “*midpoint”* (Eq. 8). The “correct” parameter-fitting solution yields a value for *iota*_*midpoint*_ such that *x*_norm._ = 1 (Eq. 10). The value of *iota* depends on three unknown parameters, x^2^/D_eff_, k_for_ and k_rev_ = k_for_ * K_D_, needed to calculate the values of *eta* and *kappa*_*f*_. To obtain these, the previously determined values of [Ab_1_], [AgAb]_equil_1_ and K_D_ were tested together with pairwise combinations of x^2^/D_eff_ (0.25 to 4000 s; n=55) and k_for_ (3E3 to 1E7 M^-1^s^-1^; n = 58) values to find the solutions for *iota*_*midpoint*_ yielding x=1, y = *f* = 0.5. While there are an infinite number of solutions that satisfy this condition, the various time courses differ from each other at other values of *f*. The predicted time course for each pair of parameters is therefore compared with the observed time course at 27 other values of *f* ranging from 0.001 to 0.95 to find the parameter pair with the minimum average error (f_observed_ – f_predicted_), and that pair are used for the next step.

In the second step, parameters near the x^2^/D_eff_ and k_for_ pair found in the previous step are iteratively re-tested across a decreasing range, using a log step scale (*i*.*e*., candidate values are more closely spaced near the center of the scale). The initial range is ± 10-fold from the central value but is gradually reduced with each cycle until a stable result is obtained or 8 cycles are completed.

The third and last step refines the candidate k_for_ and x^2^/D_eff_ pairs by sampling 841 parameter combinations (29 values of k_for_ = 3E3 to 1E6 M^-1^sec^-1^; 29 values of x^2^/D_eff_ = 0.3 to 1E4 s) distributed around the parameter pair found in the previous step, again calculating an average fractional error between the predicted and experimentally observed reaction time course values. The optimal x^2^/D_eff_ and k_for_ are those yielding the minimum error. In practice, the optimum parameter values typically change very little between the second and third step, but the third step provides data allowing the contour plot visualization described below.

A contour plot relates the negative log_10_ of the residual error values (Z axis) to each tested k_for_, x^2^/D_eff_ parameter pair (X and Y axes, respectively). The minimum residual error value corresponds to the predicted reaction conditions most consistent with the observed experimental data. Empirical testing showed that a contour drawn at 3x the minimum error value typically encompasses the range of experimental means ± 2 SD of the [AgAb] values at each time point. Interpretation of the overlapping contour plots for the 5-6 [AgAb]_equil_ and K_D_ values tested for each ROI is facilitated by “heat map” plots of the average error of the data sets, with the fractional error ranges color-coded.

To evaluate the performance of the fitting method, as reported in Table S4, x^2^/D_eff_ and k_for_ parameter fitting errors were quantified as the root mean square of the fractional difference between the observed vs “true” x^2^/D_eff_ and k_for_ values:

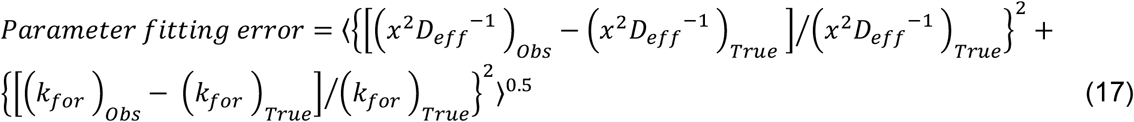

