## Supplementary figures and images for "Practical quantification of immunohistochemistry antigen concentrations and reaction-diffusion parameters"

### Supplemental movie Fig. 8

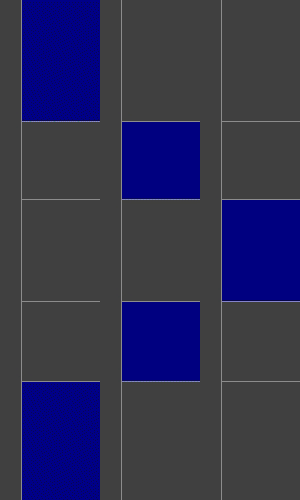
